# Phylogenetically estimated neutral rates and fitness effects of mutations to influenza proteins

**DOI:** 10.64898/2026.05.18.725477

**Authors:** Hugh K. Haddox, Angie S. Hinrichs, Chris Jennings-Shaffer, Karrington Johnson, Chelsea T. Benton, Jared G. Galloway, Jesse D. Bloom, Frederick A. Matsen

## Abstract

Influenza virus’s rapid evolution is shaped by both neutral mutation and selection. Phylogenetics can be used to study these processes, but this approach has typically only been applied to a few thousand influenza genome sequences at once. Here, we built phylogenetic trees with >100,000 influenza sequences, and then used these trees to estimate neutral rates of mutations to the virus’s genome. Neutral rates varied by up to ~100-fold among the 12 nucleotide mutation types (`A`→`C`,`A`→`G`, etc.). These rates were highly correlated among influenza, SARS-CoV-2, and HIV, though more nuanced context-dependent patterns showed marked differences between influenza and SARS-CoV-2. We also estimated fitness effects of mutations by comparing the number of times a mutation was observed to occur along the branches of a tree to the number of times we expect it to have occurred under neutrality. We estimated effects for ~33,000 nonsynonymous and ~8,000 synonymous mutations spanning all influenza proteins. This compendium of estimated effects helps map the relationship between sequence and fitness in a natural setting, including regions where synonymous mutations are under functional constraint, and for proteins with limited experimentally measured effects. We built interactive heatmaps of the estimated fitness effects to help readers explore these data (see https://matsen.group/flu-mut-rates). Altogether, this work places influenza’s mutation rates in a broader cross-viral context and deepens our understanding of how mutation and selection shape influenza evolution in nature at a site-specific level.

## Introduction

Influenza virus’s rapid evolution enables it to evade host immunity [1] and jump species barriers [2, 3]. A central goal in the field is to quantify the evolutionary forces acting on the virus. This includes both neutral mutation (i.e., the rate at which mutations arise in the absence of natural selection) and selection itself.

Quantifying a virus’s neutral mutation process is valuable because it can suggest transmission routes and underlying mutational mechanisms [4, 5, 6, 7]. It can also be used as a neutral baseline to estimate selective effects of mutations [8, 9], and to guide phylogenetic methods [10].

Quantifying selection is valuable because it can identify which sites drive immune or drug resistance [1, 11, 12, 13] or host adaptation [2, 14], and which sites are functionally constrained and thus less likely to acquire mutations that escape immunity and therapeutics [8, 15]. It can also help understand basic sequence-structure-function relationships [16, 17, 18, 19, 20]. Despite a wealth of previous work on influenza, there is still more to uncover about these processes, especially at the level of individual sites and in a natural setting where these processes are difficult to measure.

One can learn about the forces of mutation and selection at individual sites using phylogenetic trees reconstructed from hundreds of thousands or millions of viral sequences. Methods for building such trees emerged during the SARS-CoV-2 pandemic, including `UShER` [21, 22] to build parsimony-based phylogenetic trees of millions of SARS-CoV-2 sequences. Several studies have used these trees to examine SARS-CoV-2’s neutral mutation rate [4, 6, 23, 24, 25]. This rate varies dramatically among sites in the genome and is influenced by local sequence context, genomic region, and RNA structure [25]. This rate has also shifted over evolutionary time [4, 24]. Additionally, studies have used the trees to estimate fitness effects of thousands of mutations to SARS-CoV-2 proteins [8], helping to quantify selection on these proteins in a natural setting.

Here, we sought to use large-scale phylogenetics to examine neutral mutation and selection in influenza evolution. Two recent studies have used this style of approach to quantify influenza’s mutation spectrum, finding variation in mutation spectra that were suggested to be related to transmission route [5], and to estimate fitness effects of mutations to the hemagglutinin protein of human seasonal influenza with a goal of improving evolutionary forecasting for vaccine-strain selection [26]. But there are still open questions that previous studies have not fully addressed: How much does influenza’s neutral mutation rate differ between sites in the genome, and what factors cause those differences? How similar are influenza’s neutral mutation rates to those of other viruses? And what are fitness effects of mutations across all influenza proteins in a natural setting?

## Results

### Building large-scale phylogenetic trees

We built phylogenetic trees of influenza sequences and then used those trees to estimate substitution rates (Figure 1A). As input for building the trees, we used all influenza A virus sequences in the GISAID database [27] with submission dates before mid-to-late 2025 (Table S1). This corresponded to ~500,000 partial or full genome sequences, where each sequence is a consensus sequence from an infected host. Sequences came from a variety of hosts (Table S2) and subtypes (Table S3), as defined by the genotypes of the hemagglutinin (HA) and neuraminidase (NA) genes. Most came from H1N1 or H3N2, which cause human seasonal influenza.

**Figure 1:**
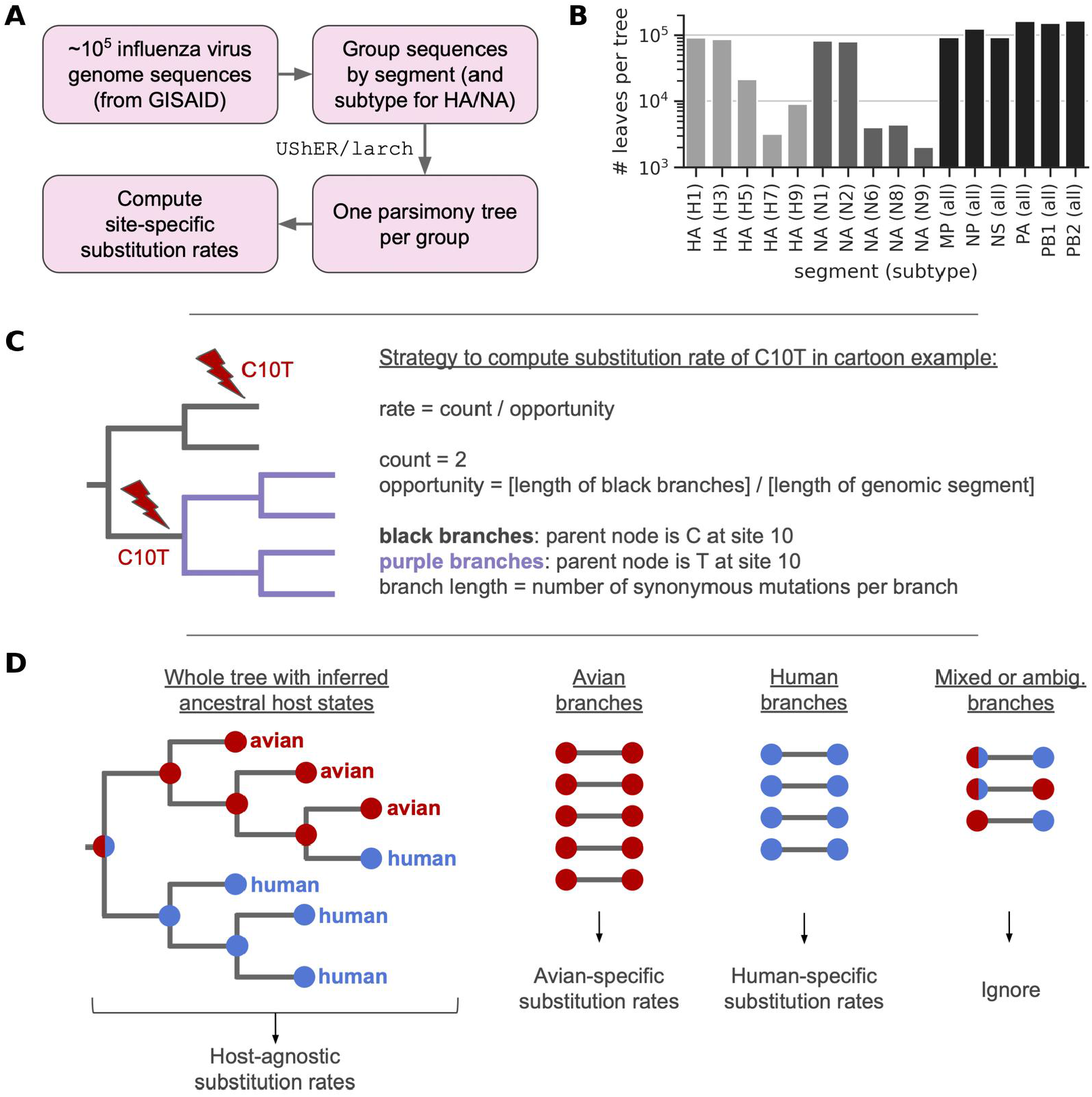
Strategy to build large-scale trees and estimate site-specific substitution rates. **(A)** Overall workflow. **(B)** Number of leaf sequences per tree for each combination of segment and subtype. We did not build trees for HA and NA subtypes that are not shown due to limited sequencing data. **(C)** Strategy to compute site-specific substitution rates illustrated using a simple hypothetical example (see *Methods* for a more formal description). **(D)** We computed host-agnostic substitution rates using trees with sequences from all hosts, using all branches passing quality-control filters (filters not depicted). We also computed host-specific substitution rates by first inferring the most parsimonious host state at each internal node of a tree, then binning quality-filtered branches by host, only considering branches where both the parent and child nodes of a branch come from a given host, and then separately computing substitution rates for each set of host-specific branches. The schematic illustrates this approach for a tree of avian and human sequences that do not cluster monophyletically by host due to the presence of a human sequence in an otherwise avian clade. The approach ignores branches with mixed host states, as well as branches with host states that could not be unambiguously assigned using the parsimony inference method.

We built parsimony-based trees using `UShER` and `larch` [28]. Because many genomes were incomplete, and because reassortment of influenza genomic segments can cause phylogenetic artifacts, we built separate trees for each segment rather than one tree for the entire genome. For the HA and NA segments, which are highly diverse between subtypes, we built separate trees for each segment and subtype. For instance, for HA, we built one tree with all HA sequences from the H1 subtype, regardless of NA subtype. We deduplicated and filtered sequences before building trees. For HA and NA, trees of highly sequenced subtypes like H1, H3, N1, and N2 had ~100,000 sequences per tree, while other subtypes had fewer sequences (Figure 1B). For the other segments, there were ~100,000 sequences per tree.

### Estimating substitution rates from trees

For each tree from above, we used the tree to estimate substitution rates at the level of individual sites in the corresponding genomic segment. We distinguish between mutation rates and substitution rates. Mutation rates reflect the neutral mutation process in the absence of selection. Substitution rates, which we estimate by counting changes along the branches of a phylogenetic tree, are shaped by both neutral mutation and selection.

We computed substitution rates as follows. First, we counted the number of times a given nucleotide mutation at a given site was observed to occur along the tree’s branches (Figure 1C). We then computed each mutation’s rate by normalizing a mutation’s observed count by its “evolutionary opportunity” on the tree (Figure 1C). We define a mutation’s evolutionary opportunity as the total number of synonymous mutations along all branches on which a mutation is possible, divided by the sequence length of the corresponding genomic segment. In the hypothetical example in Figure 1C, the C10T mutation is only possible on branches where the parent node has a C at site 10; it is not possible on other branches.

In the influenza data, values of evolutionary opportunity spanned several orders of magnitude for different mutations (Figure S1). At some sites, one nucleotide identity dominates the parent nodes across the tree. A mutation starting from that conserved nucleotide is possible on almost every branch, giving it high evolutionary opportunity. At less-conserved sites, no single nucleotide dominates the parent nodes, so for any given mutation only a fraction of branches qualify as possible. The evolutionary opportunity of any specific mutation at such a site is correspondingly lower. These differences in opportunity bias a mutation’s observed count. Normalizing a mutation’s count by its opportunity corrects for this bias in the resulting rates.

We also estimated host-specific substitution rates for human, avian, and swine hosts, ignoring other hosts due to limited data. The above trees were built using all sequences from all hosts, and the rates computed from them are host-agnostic. To infer host-specific rates, we first inferred the most parsimonious host state for each internal node in each of the above trees. For a given host, we extracted all branches for which both the parent and child nodes of a branch were inferred to have that host state, and then used those branches to infer substitution rates for that host (Figure 1D). This strategy was necessary because sequences in trees did not always cluster monophyletically by host.

Altogether, we estimated substitution rates for tens of thousands of mutations across the influenza genome. We only analyzed mutations at coding sites, due to limited sequencing coverage at non-coding sites. Below, we use data on synonymous mutations to estimate mutation rates. Then, we use these neutral rates as a baseline to estimate selection on individual synonymous, nonsynonymous, and nonsense mutations.

### Genome-wide mutation rates are similar between influenza viruses from different hosts and between influenza, SARS-CoV-2, and HIV

We estimated mutation rates using synonymous mutations. We assume that synonymous mutations have neutral effects, such that their substitution rates correspond to mutation rates [29]. Accordingly, we excluded synonymous mutations in overlapping reading frames in the PA, MP, and NS segments, unless they were synonymous in both frames. In an analysis described below, we detected strong purifying selection on synonymous mutations in regions with packaging signals at segment termini and regions near splice sites for the M2 and NEP coding sequences. So, we excluded these regions when estimating mutation rates, as well.

To start, we estimated the genome-wide mutation rate of each of the 12 nucleotide mutation types. We did so by averaging synonymous substitution rates across sites. The resulting values showed a wide spectrum of rates that varied by ~100-fold between the mutation types with the highest rates (`G`→`A` and `C→T`) and those with the lowest rates (`C→G` and `G→C`) (Figure 2A). Host-specific rate spectra were highly similar to one another (Figure 2A). Below, we analyze host-agnostic rates, unless otherwise noted.

**Figure 2:**
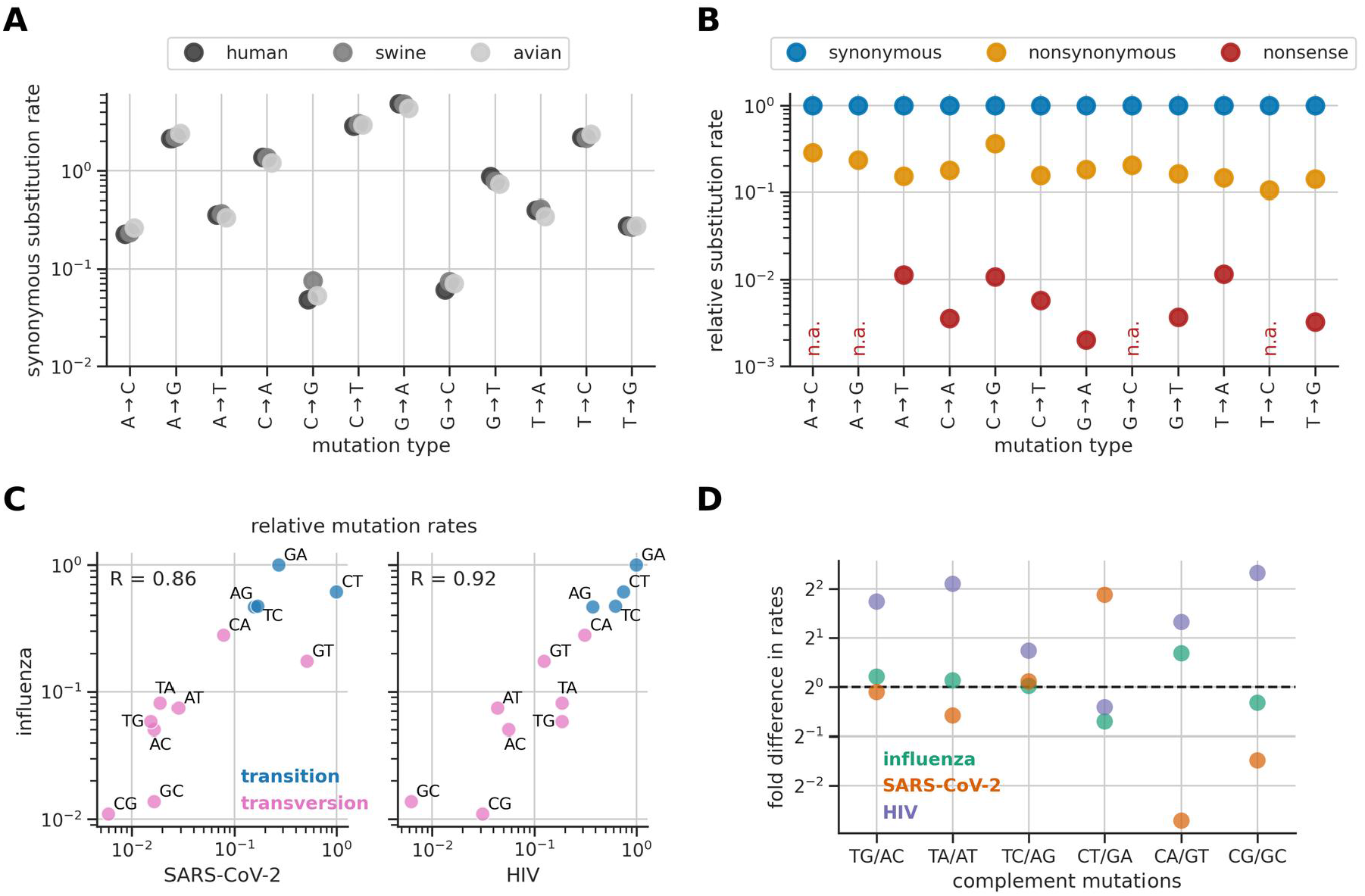
Genome-wide synonymous substitution rates. **(A)** Rates of synonymous mutations vary widely between mutation types. This spectrum of rates is similar between hosts. **(B)** Rates of nonsynonymous and nonsense mutations are lower or much lower than rates of synonymous mutations. Rates are expressed relative to the synonymous rate for the same mutation type (“n.a.” indicates mutation types that never lead to nonsense mutations). **(C)** Mutation rates are correlated between influenza, SARS-CoV-2 [25], and HIV [30]. Rates for each virus are normalized so that the mutation with the highest rate has a value of one. **(D)** Mutation rates are most similar between pairs of complement mutation types (fold difference closest to one) for influenza. Panels B-D use the host-agnostic rates we estimated for influenza virus.

To assess data quality, we also computed average substitution rates of non-synonymous and nonsense mutations. These rates were ~5-fold and ~100-fold lower than rates of synonymous mutations, respectively (Figure 2B), consistent with strong purifying selection. This result suggests that artificial sequencing errors do not substantially inflate synonymous rates, as sequencing errors are not expected to be selectively purged.

Next, we compared the mutation rates we estimated for influenza to those of SARS-CoV-2 and HIV, as estimated in previous studies [25, 30]. Rates were remarkably similar between viruses (Figure 2C). This is despite the fact that they use substantially different replication machinery. Influenza and SARS-CoV-2 both use RNA-dependent RNA polymerases, but they are highly diverged [31] and the SARS-CoV-2 machinery includes a proofreading mechanism [32, 33]. HIV uses a reverse transcriptase, along with a host DNA-dependent RNA polymerase. The similarity in rates between viruses suggests that they share at least some common underlying mutational mechanisms. For instance, transition mutations tend to occur at higher rates than transversions for all three viruses (Figure 2C), suggesting common mechanisms related to chemical similarities between transitions. However, the transition bias only explains some variability in rates, suggesting other common mechanisms are also at play.

Interestingly, influenza shows substantially higher symmetry in rates between complement mutation types (Figure 2D). The rates from Figure 2 are all computed relative to a specific strand. If rates are similar between complement mutation types, that indicates the mutation process is symmetrical between strands. If they are not similar, that indicates asymmetry. Both SARS-CoV-2 and HIV show substantial asymmetry for at least one mutation type, while influenza showed considerably more symmetry. Thus, although rates are highly correlated between viruses (Figure 2C), we find clear differences in degree of symmetry between strands.

### Site-specific mutation rates depend on local sequence context, with context-dependent patterns differing between influenza and SARS-CoV-2

Next, we explored how much mutation rates varied between sites. As above, we estimated mutation rates using substitution rates of synonymous mutations, excluding sites in regions of functional constraint. We found substantial variability in site-specific mutation rates (Figure 3A), though less than the level previously observed for SARS-CoV-2 [25] (Figure S3).

**Figure 3:**
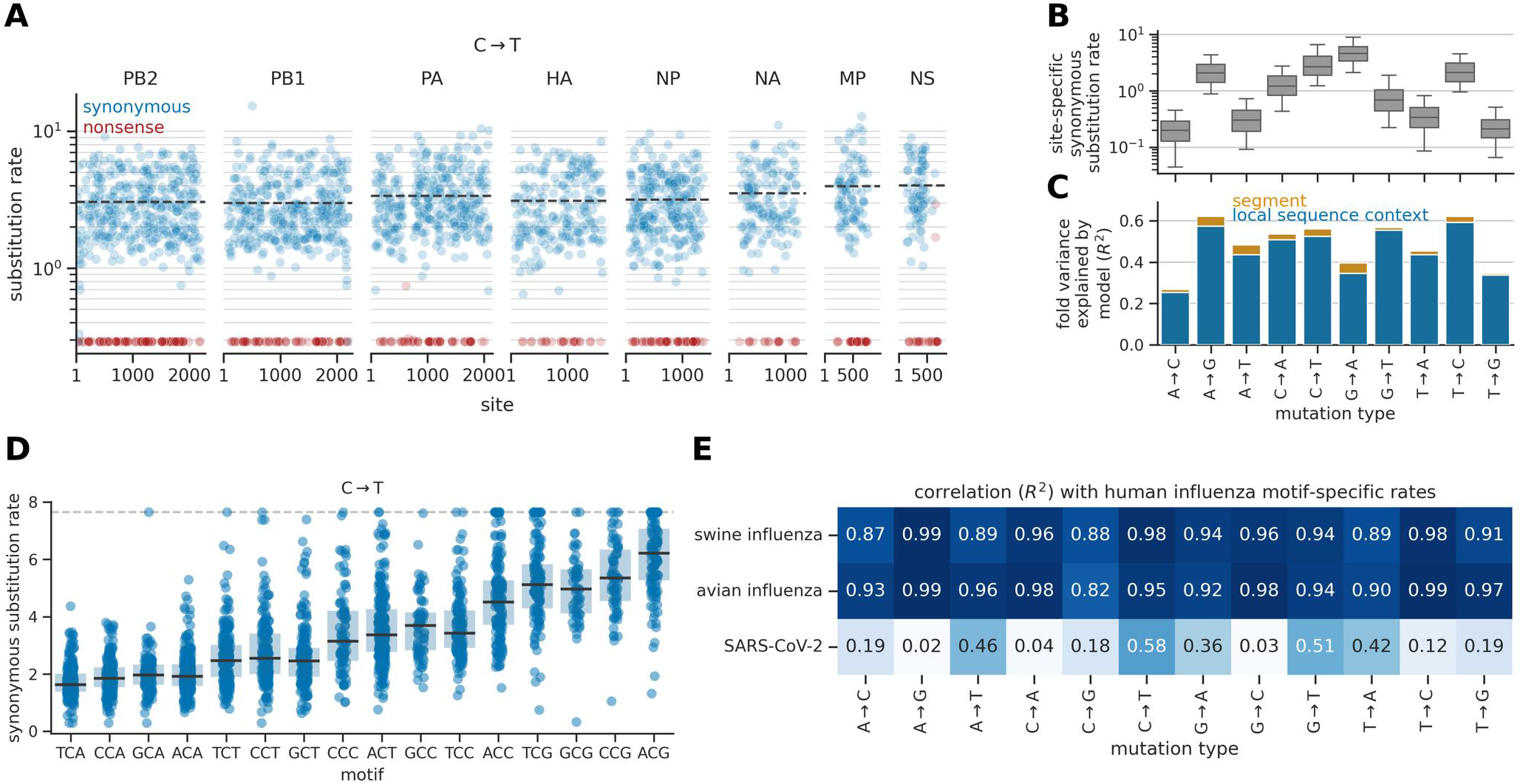
Patterns of site-specific synonymous substitution rates. **(A)** Site-specific rates of synonymous `C→T` mutations at coding sites along the genome, clipped at a lower limit of detection (see *Methods*). Sites are numbered in context of positive-sense strands, ranging from the first coding site to the last coding site (we ignored non-coding sites). Blue dots show synonymous mutations and red dots show nonsense mutations. This and other panels only include synonymous mutations used to estimate mutation rates, which excludes certain mutations in overlapping reading frames and the regions of functional constraint listed above. Dashed gray lines show average synonymous rates for each segment. For HA and NA, we only show rates for the H1 and N1 subtypes, respectively. **(B)** Distribution of site-specific synonymous rates for each mutation type (there were not enough data to estimate site-specific rates for `C→G` and `G→C` mutations). **(C)** The fold variance in site-specific synonymous substitution rates explained by the neutral mutation model, quantified by the correlation between observed and predicted log rates (*R*^2^). The blue bar shows the correlation for a model trained only on local sequence context. The orange bar shows the increase in correlation for a model trained on both local sequence context and genomic segment. **(D)** Site-specific synonymous `C→T` substitution rates as a function of local sequence context, defined by a site’s 3-mer motif, with a ceiling on plotted points at the 98th percentile of the overall distribution (horizontal dashed line). **(E)** The correlation in motif-specific synonymous substitution rates between human influenza and either swine influenza, avian influenza, or SARS-CoV-2 [25].

We sought to identify factors that help explain this variability. Motivated by previous work on SARS-CoV-2 [6, 25], we examined three main factors: local sequence context, genomic region, and RNA structure. For influenza, the strongest determinant was local sequence context, as defined by the 3-mer nucleotide motif centered on the mutated site. For example, the median rate of `C→T` mutations varied by about four-fold between motifs (Figure 3D). Most other mutation types showed a similar range of variation, though a few showed a higher range (Figure S4). A log-linear model trained to predict a site’s synonymous substitution rate based on its local sequence context can explain ~20-60% of the fold-variation in rates between sites, depending on mutation type (Figure 3C).

Some variation in rates between motifs might be caused by host restriction factors. As in Ruis et al. [5], we saw that mutations that ablate a CG dinucleotide motif tended to have elevated rates compared to mutations in other contexts (Figure 3D; Figure S4), consistent with selection to evade ZAP, which targets such motifs [34]. We also searched for signatures of APOBEC, which can introduce C→T mutations in viral genomes [35, 36, 37]. One motif with a relatively high C→T rate is a known target of APOBEC3A (TCG; Figure 3D) [37], though the high rate at this motif could also be explained by selection to evade ZAP.

Context-dependent patterns were highly similar between influenza viruses from human, swine, and avian hosts, but strikingly different between influenza and SARS-CoV-2 (Figure 3E). This result suggests that context-dependent effects largely arise through different underlying mechanisms for these two viruses.

Unlike SARS-CoV-2, we did not find evidence that rates substantially depend on genomic region or RNA secondary structure. Adding a site’s segment as a predictor variable in the log-linear model did not substantially increase the model’s ability to predict synonymous substitution rates (Figure 3C). Addition-ally, site-specific synonymous substitution rates were only weakly correlated with high-throughput experimental data quantifying a site’s tendency to be paired in the RNA structure of the viral genome (Figure S5) [38].

### Regions of constraint on synonymous mutations

Next, we searched for regions in the genome where synonymous mutations are under purifying selection. To do so, we computed fitness effects of individual synonymous nucleotide mutations using the approach from Bloom and Neher [8]. As above, we quantified the count and evolutionary opportunity of each mutation. We call this count an “actual count”, as it corresponds to the actual number of times a mutation was observed. For each mutation, we also computed an “expected count” that corresponds to the number of times we would have expected to see the mutation under neutrality. We did so by multiplying a mutation’s evolutionary opportunity by its neutral rate estimated by the segment-aware neutral model from above (for C→G and G→C, which are not included in the neutral model, we used mean synonymous substitution rates for a given motif).

We then estimated a mutation’s fitness effect as Δ*f*_*m*_ = log(*a*_*m*_ /*e*_*m*_), where *a*_*m*_ and *e*_*m*_ are a mutation’s actual and expected counts, respectively. We added a pseudocount of 0.5 to both the numerator and denominator to avoid division by zero. If the actual and expected counts are the same, a mutation is inferred to have a neutral effect equal to zero. If the actual count is greater than the expected count, a mutation is inferred to have a positive (beneficial) effect, and vice versa if the actual count is less than the expected count. To reduce the impact of noise, we only analyzed mutations with at least ten actual or expected counts. As above, we excluded synonymous mutations in overlapping reading frames in the PA, MP, and NS segments, unless they were synonymous in both frames.

Examining effects of synonymous mutations across the genome, we identified several regions where these effects were substantially deleterious, indicating functional constraint (Figure 4A). Most such regions occurred at segment termini, which contain packaging signals for incorporation of genomic segments into budding viral particles (see the red shaded regions) [39, 40, 41, 42, 43, 44, 45, 46, 47]. Disruption of packaging may explain the deleterious fitness effects. These results build on previous studies by helping identify specific sites where synonymous mutations putatively disrupt packaging signals, such as the clusters of sites highlighted in Figure 4B.

**Figure 4:**
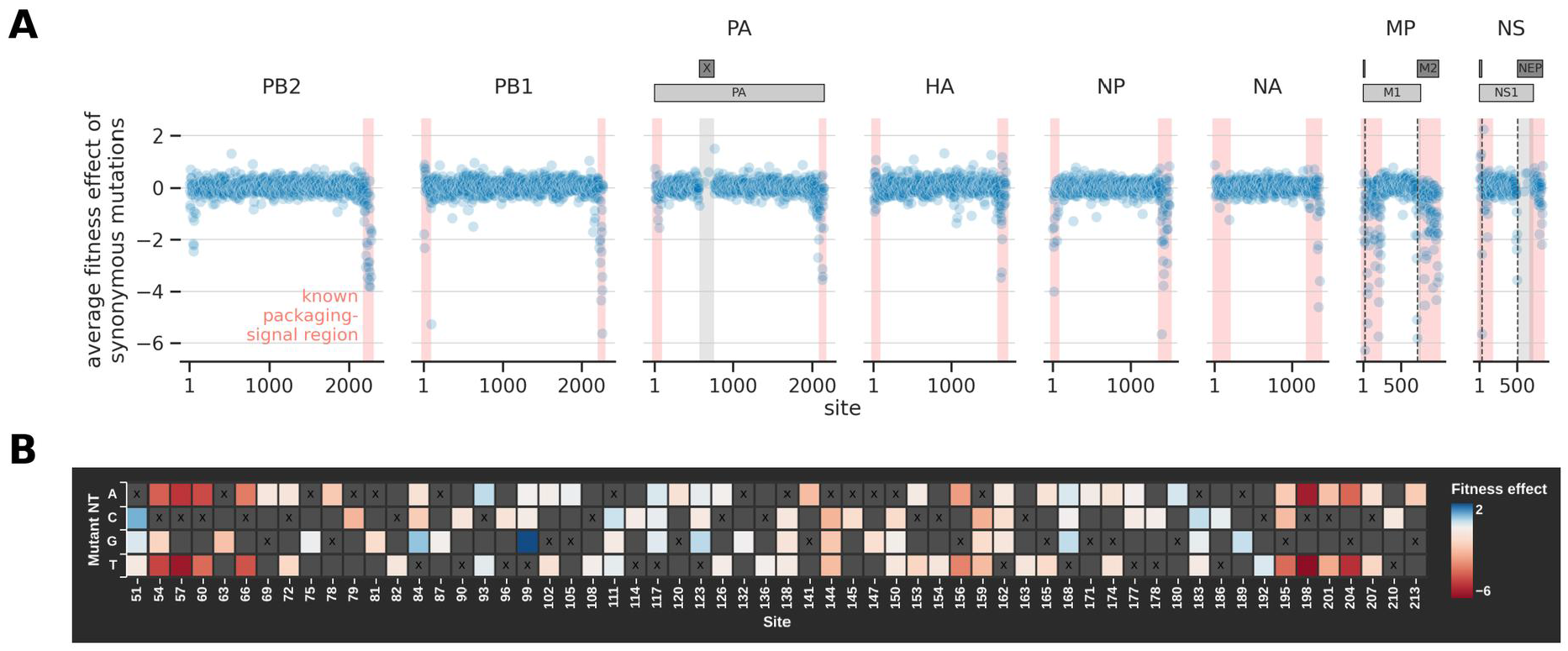
Regions of constraint on synonymous mutations. **(A)** Average per-site fitness effects of synonymous mutations across the genome (plot widths scaled by segment length). Sites are numbered in context of positive-sense strands, ranging from the first coding site to the last coding site (we ignored non-coding sites). Shaded red areas at segment termini correspond to regions experimentally identified to have packaging signals, as summarized in Li et al. [39]. The shaded gray areas show regions with overlapping reading frames (PA and X in the PA segment; M1 and M2 in the MP segment; NS1 and NEP in the NS segment). The horizontal bars above the plots of these segments show the location of these reading frames, including multiple exons for both M2 and NEP. The vertical dashed lines show the location of exon/intron boundaries involved in splicing of M2 or NEP. The shaded gray areas show regions of overlap between M1 and M2 or NS1 and NEP, where we excluded synonymous mutations unless they were synonymous in both frames. For HA and NA, we only show effects for the H1 and N1 subtypes, respectively. **(B)** A heatmap of fitness effects of synonymous nucleotide mutations near the start of the MP segment, showing multiple regions of constraint. See https://matsen.group/flu-mut-rates/nt for interactive heatmaps of fitness effects of nucleotide mutations across all coding sites of all segments.

In the MP and NS segments, synonymous mutations also had deleterious effects near exon/intron boundaries associated with splicing of the M2 and NEP coding sequences, respectively (see the vertical dashed lines in Figure 4A). For three of the boundaries, we detected constraint at sites immediately at the boundary; for the fourth, we detected constraint at the nearest site with an estimated effect (Figure S6).

To further explore these data, see https://matsen.group/flu-mut-rates/nt for interactive heatmaps of fitness effects across all sites and segments.

### Estimated fitness effects of amino-acid mutations

Next, we estimated fitness effects of individual amino-acid mutations to each protein in the genome. We used the same approach as above, but first aggregated counts over all nucleotide mutations that resulted in the same amino-acid mutation before computing that mutation’s fitness effect.

Broadly speaking, the fitness effects followed expected trends (Figure 5A). Most synonymous mutations had effects near zero. Nonsynonymous mutations had a wide range of effects, mostly deleterious. Most nonsense mutations had very deleterious effects.

**Figure 5:**
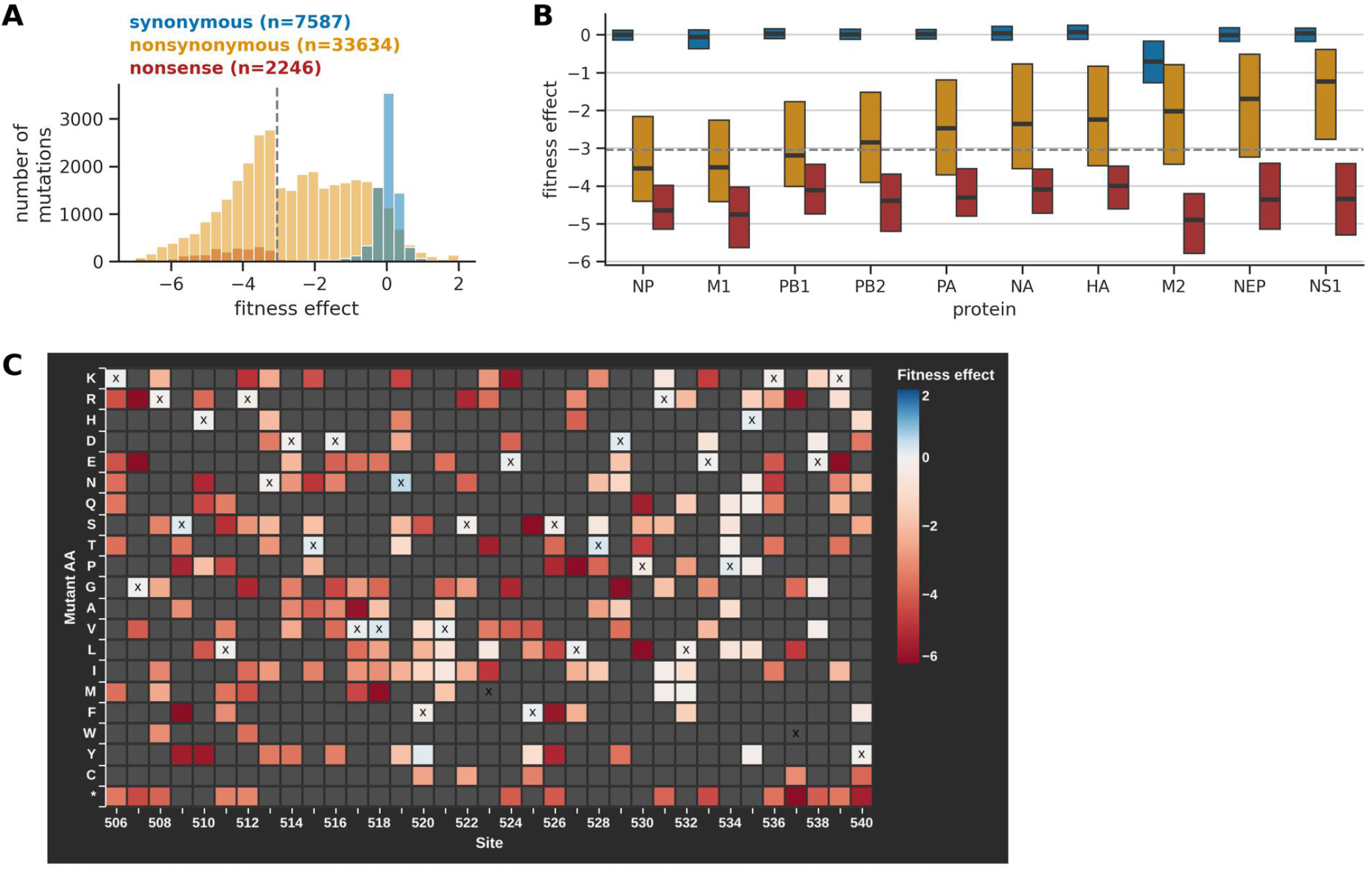
Estimated fitness effects of mutations. **(A)** Distribution of fitness effects for synonymous, nonsynonymous, and nonsense mutations. The legend reports the number of mutations per category. The vertical dashed line shows a lower limit of detection corresponding to the fitness effect of a mutation with an actual count of zero and an expected count of ten. A mutation’s effect can be more negative than this line only if its expected count is greater than ten. **(B)** Distributions of fitness effects by protein, with one box plot per mutation class: synonymous, nonsynonymous, and nonsense, using the same color scheme as panel A. The horizontal dashed line shows the same limit of detection as panel A. **(C)** A heatmap of fitness effects of mutations to a short functionally constrained region of PA not previously characterized by DMS. See https://matsen.group/flu-mut-rates for complete interactive heatmaps of mutational effects to all proteins.

Different proteins had different levels of tolerance to nonsynonymous mutations (Figure 5B). The median fitness effect of nonsynonymous mutations ranged from −3.5 for NP to −1.1 for NS1. The two proteins with the lowest mutational tolerance (NP and M1) both form oligomers that interact with numerous nucleic-acid or protein partners [48]. Their low mutational tolerance could arise from these various structural and functional constraints. Despite these differences, all proteins were highly intolerant to nonsense mutations (Figure 5B), indicating all are important for viral fitness in nature.

M2 stands out as having the lowest tolerance to synonymous mutations (Figure 5B). As above, in overlapping reading frames in the MP and NS segments, we only classified mutations as synonymous if they were synonymous in both frames. So, the constraint we observe is not due to overlapping reading frames. Rather, the constraint could be because the M2 coding sequence overlaps extensively with regions with known packaging signals (Figure 4A).

Altogether, we estimated fitness effects for ~33,000 nonsynonymous and ~ 8,000 synonymous mutations across the genome. In the case of HA and NA, this included separately estimating effects for different subtypes. These data provide a rich catalog of mutational effects in a natural setting, including mutations that have not been previously experimentally characterized. See https://matsen.group/flu-mut-rates for interactive heatmaps of fitness effects. Figure 5C shows a snapshot of a heatmap for a functionally constrained region of the PA protein.

A caveat is that we estimated one effect per mutation for each input tree. Thus, our estimates do not capture shifts in mutational effects over evolutionary time along a given tree, which can occur due to epistasis or changes in external selective pressures like host immunity [16, 49, 50].

### Fitness effects of amino-acid mutations are robust to sub-sampling and correlate with experiments

The estimated fitness effects are subject to statistical sampling noise due to limited mutational counts in the available sequences. To estimate levels of noise, we randomly divided the branches of the phylogenetic trees into two equal subsets, and then separately computed fitness effects from each subset. Effects were well correlated between subsets (Figure 6A), suggesting the effects are robust to subsampling, despite some noise.

**Figure 6:**
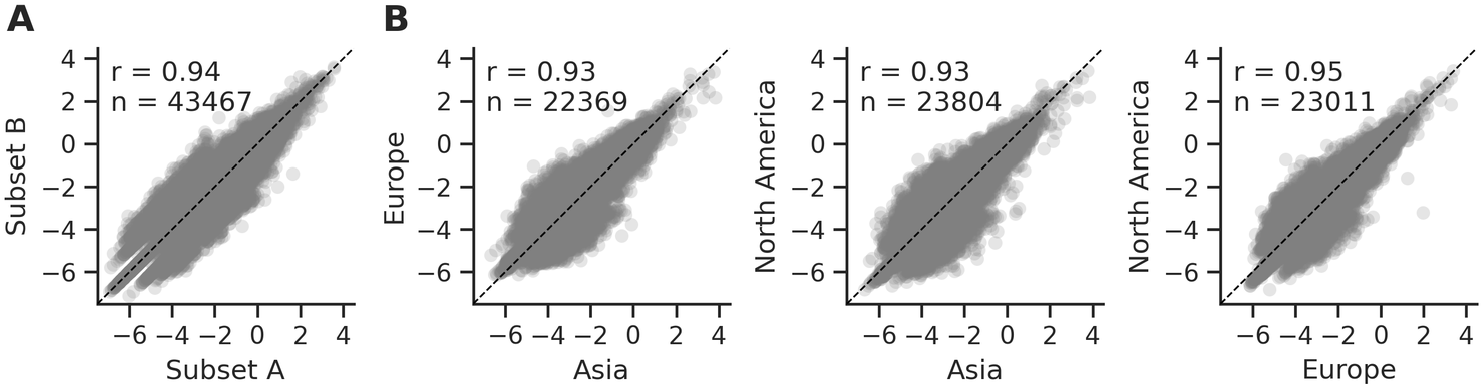
Estimated fitness effects of amino-acid mutations are robust to subsampling. **(A)** Fitness effects are correlated when randomly dividing the input branches into two equal subsets (A and B) and then separately estimating effects from each subset. The plot shows all mutations from Figure 5A, which each have at least ten actual or expected counts when summed across the two subsets. **(B)** Fitness effects are correlated when extracting subtrees with all leaf nodes from a given region, and then separately estimating effects from each subtree. Each plot compares two regions and shows all mutations with at least ten actual or expected counts in each of the two regions, so as to focus on mutations with sufficient data to accurately estimate effects in each region. *r* is the Pearson correlation coefficient and *n* is the number of plotted mutations.

To search for signal of bioinformatic artifacts, we extracted subtrees with all leaf nodes from either Asia, Europe, or North America, and then separately computed fitness effects from each subtree. Effects were well correlated between subtrees (Figure 6B), suggesting regional differences in sequencing and bioinformatic workflows do not substantially bias the results.

To help validate the estimated fitness effects, we also compared them to effects experimentally measured by deep mutational scanning (DMS). Most influenza proteins have been characterized by DMS. We focused on DMS experiments that select for a protein to fold and perform one or more basic functions for viral replication in cell culture or a model organism [12, 13, 14, 16, 17, 18, 19]. We did not include a recent DMS of the NEP protein [20], since the selection in that experiment did not include constraint due to overlapping reading frames between NEP and NS1.

For most experiments, there is an intermediate-to-high correlation between our estimated fitness effects and DMS-measured effects among mutations with data from both approaches (Figure 7). The correlation is highest for HA, with a Pearson correlation coefficient of 0.82. These results suggest our fitness effects estimated in a natural setting often capture basic functional constraints measured in defined experimental conditions.

**Figure 7:**
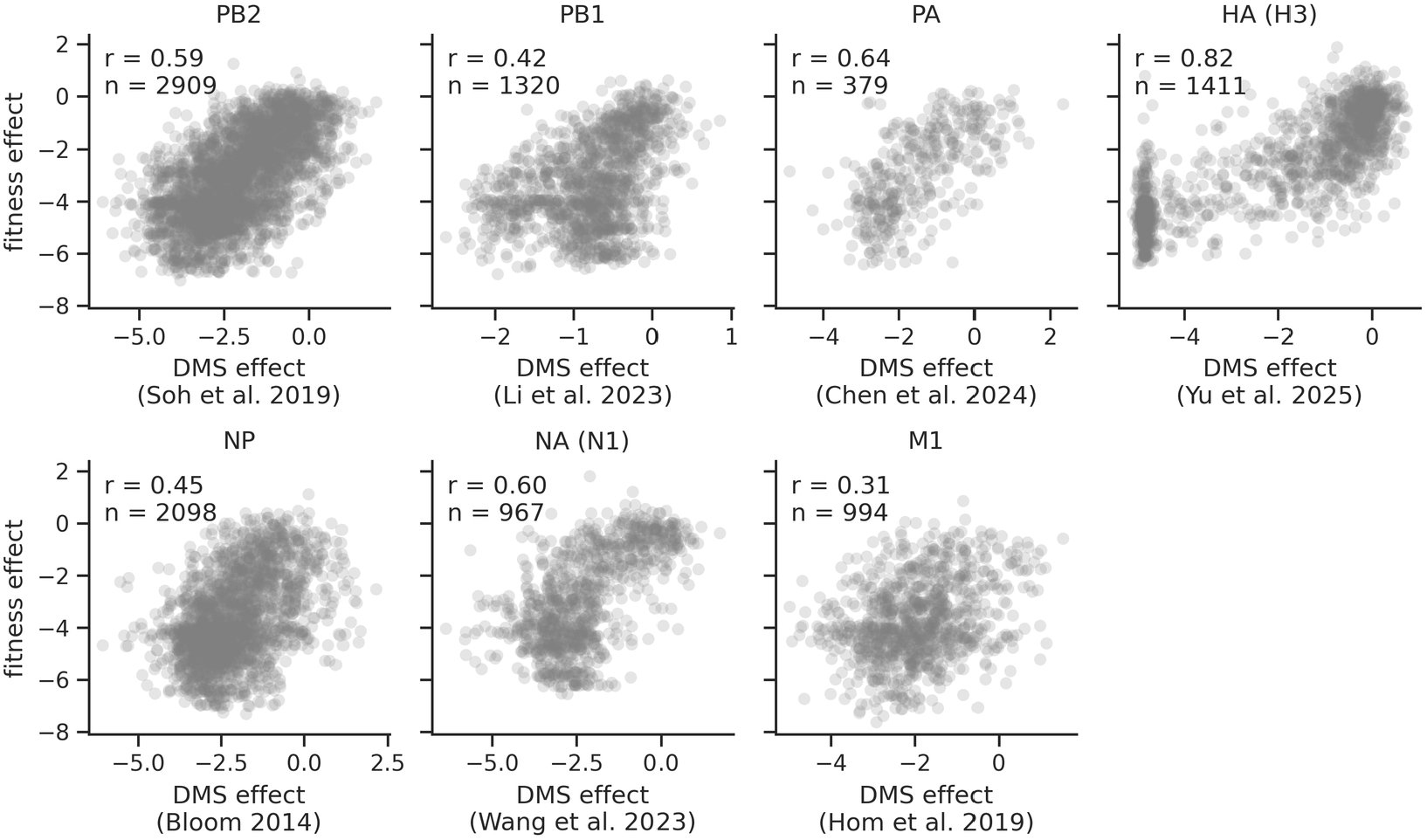
Estimated mutational effects correlate with deep mutational scanning data. Each plot shows the correlation between mutational fitness effects estimated here and effects measured by DMS for a given protein (*r* is the Pearson correlation coefficient and n is the number of plotted mutations with effects estimated both here and in the DMS experiment). X-axis labels give citations for the DMS experiments. These plots only show data for mutations with ≥ 25 actual or expected counts.

For a few experiments, the correlation is low, with the lowest value being 0.31 for M1. Differences between fitness effects and DMS effects could be due to a number of factors, including: noise in our estimates or the experiment, and differences in selection between lab and nature.

## Discussion

We used large-scale phylogenetics with hundreds of thousands of influenza sequences to examine neutral mutation and selection in influenza evolution at the level of individual sites in a natural setting.

On the side of neutral mutation, we compared influenza’s mutation spectrum to that of SARS-CoV-2 and HIV. We found remarkable concordance between the overall rates of the 12 nucleotide mutation types between viruses. At the same time, we found clear differences in more nuanced context-dependent rates between influenza and SARS-CoV-2 (the comparison to HIV was not possible for lack of prior estimates). These results point to common mutational mechanisms underlying the differences in rates between the 12 mutation types, which span two orders of magnitude, but different mechanisms underlying context-dependent rates, which often span less than an order of magnitude for a given mutation type.

Compared to SARS-CoV-2, influenza’s neutral spectrum was perhaps less “surprising” overall: it is more symmetrical between strands and less variable between sites. A simple hypothesis to explain the greater symmetry is that most influenza mutations come from the viral polymerase and that polymerase errors occur equally on both strands. A simple hypothesis to explain SARS-CoV-2‘s higher variability is that its replication machinery involves a proof-reading mechanism [32, 33], while influenza’s machinery does not. It is possible the efficacy of proofreading is context dependent, which could amplify pre-existing biases.

Having quantified influenza’s neutral mutation spectrum, we then used it as a baseline to estimate fitness effects of mutations. We estimated effects for ~ 33,000 nonsynonymous mutations and ~8,000 synonymous mutations across all influenza proteins.

This compendium of estimated effects builds on previous work in three main ways. First, we estimated effects of amino-acid mutations that have not been previously experimentally characterized by DMS. No DMS experiments have been performed on M2 or NS1, and the DMS experiment on PA only covered the first 240 of 716 amino-acid sites [13]. Our estimates help map the relationship between sequence and function for these proteins.

Second, the estimated effects of synonymous mutations help to more comprehensively map functional constraint on these mutations across the genome. Specifically, we detected constraint in regions known to be important for packaging of genomic segments and splicing. In the past, packaging signals have been identified by making large deletions or by introducing clusters of mutations in packaging sequences, often in genetic constructs with artificial reporter genes [39, 40, 41, 42, 43, 44, 45, 46, 47]. The estimated effects we report help map specific sites of putative importance for packaging in a natural setting.

Third, we estimated effects of mutations in a natural setting. Experiments measure effects under controlled lab conditions, which are useful for isolating specific selective pressures, but do not fully capture the complex set of selective pressures in nature. Comparison of the fitness effects we estimated here and ones measured in the lab could be used to identify regions where selection differs between these settings.

Previous studies have identified differences in the nucleotide composition of influenza viruses between human and avian hosts [51, 52, 53], including a large decrease in the number of CG dinucleotide motifs during the evolution of H1N1 influenza in humans between 1918 to 2009, following its spillover from the avian reservoir [52]. These studies have proposed a number of mechanisms to explain these observations, including host-specific differences in both mutation and selection. In the mutation rates we estimated, C→T mutations have one of the highest rates (Figure 2A). For this and other mutation types, their rates are often higher in contexts that ablate CG motifs (Figure S4). These high rates could help explain why CG motifs were depleted in H1N1 during evolution in humans. However, in our estimates, these rates are equally high in human and avian hosts (Figure S7). So, our estimates do not explain why CG motifs would have been specifically depleted in humans. Possibly, the depletion is driven by selective pressures that are not apparent in the site-averaged synonymous substitution rates we examined. Or possibly, it is driven by mutational processes not captured in our analysis due to methodological limitations, such as the ones described next.

Our study has several limitations. First, in analyzing host-specific mutation rates, we only considered three groups: human, swine, and avian. In contrast, a recent study by Ruis et al. [5] partitioned avian hosts into finer groups, finding modest but statistically significant shifts in rates between groups. Our coarser grouping did not allow for as nuanced an analysis, though we build on their study by comparing mutation rates across viruses and by examining site-specific mutation rates in greater detail. Second, our estimated mutation rates and fitness effects are averaged over many sequences. For instance, the estimates for H3 HA were derived from pooling all available H3 sequences. However, mutational effects can change over evolutionary time due to epistasis and changes in external selective pressures, such as host population immunity [16, 49, 50]. Mutation rates can also change abruptly during viral evolution [4, 24]. Our estimates do not capture such changes along a given tree of sequences. In the future, our approach could be updated to compute mutation rates and fitness effects for sequences grouped by clades [8, 26] or windows in time [54].

Overall, this work adds to our knowledge of mutation and selection on influenza in nature at a site-specific level, laying the foundation for future comparisons to other viruses and between variants of influenza over evolutionary time.

## Methods

### Data and code availability

See https://github.com/matsengrp/flu-usher for the code we used to build phylogenetic trees (see commit `0c5a4b8`). Table S1 provides GISAID EPISET IDs for all sequences used to build trees. See https://github.com/matsengrp/flu-mut-rates for the code we used to compute substitution rates and fitness effects from the above trees (see commit `e24f7d0`). This repository also includes key result files (paths to specific files are described below and in the repository’s `README.md` file).

### Building large-scale phylogenies

First, we downloaded all influenza A virus nucleotide sequences and associated metadata from GISAID [27] (https://gisaid.org/) with submission dates before mid-to-late 2025 (Table S1). We grouped sequences by genomic segment. For internal segments (PB2, PB1, PA, NP, MP, NS), we combined all subtypes into one group for analysis. For HA and NA segments, we further divided the data by subtype. In total, we built separate trees for five HA subtypes (H1, H3, H5, H7, H9) and five NA subtypes (N1, N2, N6, N8, N9), ignoring other subtypes due to limited data.

For each group of sequences from above, we aligned the sequences to a group-specific reference sequence we downloaded from NCBI (https://www.ncbi.nlm.nih.gov/; Table S4). We chose common reference strains, avoiding ones with ambiguous nucleotides (for H1, A/California/07/2009 had ambiguous nucleotides in the HA sequence, so we chose a closely related sequence). We did so using `nextclade` [55] (https://clades.nextstrain.org/), which iterates over each input sequence, performs a codon-aware pairwise alignment to the reference, and strips all insertions relative to the reference, such that all output sequences are the same length as the reference (including gap characters). Next, we curated the alignment in the following ways. We trimmed non-coding sites from the alignment (due to low sequencing coverage). We excluded sequences if: i) their coding sequences (unaligned) were not a multiple of three and did not start with a start codon and end with a stop codon; ii) they had ambigu-ous nucleotides or *>*3% gap characters; or iii) they were a duplicate of another sequence (keeping one entry per duplicate). Between ~25-50% of sequences passed these filters, depending on the segment and subtype (Figure S8).

Next, for each alignment, we built a parsimony-based phylogenetic tree using a combination of `UShER` [21, 22] and larch [28], where `UShER` was first used to generate initial trees, and `larch` was then used to find more parsimonious trees. Specifically, for a given input alignment, we created ten versions of the alignment where we randomized the order of the sequences (keeping the first sequence in the same position, as this is the designated reference/root sequence). For each of the ten alignments, we used `UShER` to build a phylogenetic tree, resulting in ten trees. These trees differed in topology and parsimony score due to stochasticity and lack of convergence in the search algorithm. We used randomly ordered alignments as inputs in an effort to promote greater topological diversity. Then, we used `larch` to merge the ten trees into a single directed acyclic graph (DAG), which is a compact way to represent many trees in one object. The larch DAG is an efficient way to search tree space for more parsimonious tree topologies. That is because the DAG made from the ten input trees actually represents a much larger number of trees, including all possible trees that can be made via combinations of the input trees. This number can be quite large. For instance, the DAG made from the ten H1 HA input trees represented ~ 10^451^ possible trees. After merging the input trees into a single DAG, we then used larch to extract a maximally parsimonious tree from the DAG. The extracted tree was more parsimonious than any of the ten input trees, sometimes by hundreds of mutations (Figure S9).

After extracting a maximally parsimonious tree from the DAG, we then rerooted certain trees using an avian sequence with a relatively early collection date (Table S5). The reference sequences from above were convenient for numbering purposes, but were not always biologically realistic roots. The early avian sequences provided more realistic roots.

Finally, we used `taxonium` [56] to create interactive visualizations of trees.

Trees are available upon request to anyone with EpiFlu GISAID access creden-tials.

### Inference of ancestral host states

The GISAID metadata reports the host associated with each sequence. We mapped hosts into one of five categories: human, avian, swine, bovine, or other (Figure S2). For each of the final rerooted trees from above, we used PastML [57] with the DOWNPASS maximum-parsimony algorithm to infer the most parsimonious host state of each internal node in the tree based on the known states of the leaf sequences.

### Site numbering scheme

We computed rates and fitness effects of mutations at the level of individual nucleotide sites. In doing so, we numbered sites in context of the positive-sense strand of each segment, ranging from the first to the last coding site of each segment. We excluded non-coding sites due to limited sequencing coverage. For a given tree, the site numbering is relative to the reference sequence used to build the tree (Table S4). We determined the location of coding regions in a reference sequence using associated gene annotations from NCBI.

### Computing substitution rates

We computed site-specific substitution rates using the following strategy. Below, we use 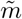 to denote a mutation type (e.g., C→T) and *i* to denote a site, and we write 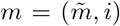 as shorthand for a mutation at a specific site, matching the notation used in the *Results*. Let 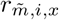 be the substitution rate of mutation type 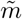 at site *i* (in the aligned sequences from a given tree) in local sequence context *x* (defined by the nucleotide identities of the sites to the left and right of site *i*). For instance, in Figure 1C, 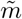 is C→T and *i* is site 10. For simplicity, that example did not include local sequence context, but an example context is A<u>C</u>A, where the underlined C is the site being mutated. We computed the substitution rate as:

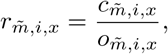

where 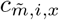 is the mutation’s count and 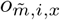 is the mutation’s evolutionary opportunity (as Figure 1C describes using informal notation). Its count is simply the number of times the mutation was observed to have occurred along the branches of the tree. Its evolutionary opportunity is defined as:

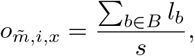

where *B* is the set of branches along which the mutation is possible, *l*_*b*_ is the length of branch *b* (defined as the total number of synonymous mutations on that branch), and *s* is the length of the alignment (after trimming non-coding sites). A mutation is possible on a branch if the mutation goes away from the branch’s parent-node sequence. For instance, in the cartoon example from Figure 1C, the C→T mutation at site 10 is possible on branches where the parent node has a C at site 10, but not possible on branches where the parent node has a T at site 10. Normalizing counts by evolutionary opportunity helps make the resulting rates comparable between mutations, as mutations span a wide range of opportunities (Figure S1). Normalizing evolutionary opportunity by alignment length helps make rates comparable between segments. Quantifying branch length using synonymous mutations helps reduce biases between segments related to differences in levels of tolerance to nonsynonymous mutations. Figure 3A/B show 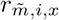 values.

We computed genome-wide substitution rates by taking a weighted average of rates across sites and contexts, with weights proportional to a mutation’s evo-lutionary opportunity. Specifically, for a specific mutation type 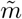, we computed the weight of that mutation type at a specific site and context as:

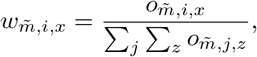

where the denominator sums over all sites *j* and contexts *z* for a fixed 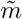. We then computed the genome-wide rate of that mutation type as:

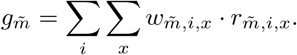

Figure 2 shows 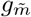values. For mutation type 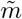in sequence context *x*, we computed its genome-wide rate as:

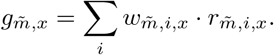

Figure 3D was computed using 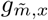 values. We used the above approach, rather than averaging 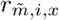 values across sites, since rates of G→C and C→G mutations were too low to estimate at the level of individual sites, but the above approach allowed us to estimate them at the genome level.

In quantifying mutation counts and evolutionary opportunity, we ignored branches that failed either of the following quality-control filters: i) there were more than four nucleotide mutations on the branch, ii) two nucleotide mutations on the branch occurred in the same codon. These filters mirror ones used in similar work on SARS-CoV-2 [8]. The first filter excludes long branches, which have the potential to be problematic in our analysis for the following reasons: parsimony-based tree-building methods can struggle to correctly place long branches in trees, and long branch lengths could arise from an abundance of artificial sequencing errors. The aim of the second filter was also to remove sequences with potential sequencing errors, as most evolution along short branches is expected to occur via single-nucleotide codon mutations. The vast majority of ignored branches were filtered out by the first filter.

We computed host-agnostic substitution rates using all quality-filtered branches from trees of sequences from all hosts. We separately computed host-specific substitution rates. For a given host, we did so using all quality-filtered branches where both the parent and child node of a branch were inferred to come from that host.

We excluded mutations from our analysis if they had low evolutionary opportunity, since lower opportunities make rates more difficult to accurately estimate. Figure S1 shows the distribution of 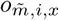 values across all mutations from a given tree. For estimating 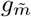 and 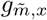 values, we excluded mutations with 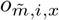 values less than 0.1. For estimating 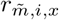 values, we excluded mutations with 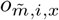 values below a mutation type-specific threshold. Specifically, for a given mutation type, we used the mutation type’s genome-wide rate to estimate the evolutionary opportunity that is expected to result in an average of ten counts per site: 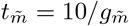. We excluded mutations for which 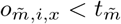. We confirmed that mutations with 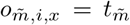 had an average of about ten counts per site, and that nearly all such mutations had at least one and often more than one count (Figure S10). This allowed us to focus on mutations with high enough counts to estimate site-specific substitution rates. No G→C or C→G mutations passed this threshold, so we were not able to estimate site-specific rates for these mutation types.

We clipped site-specific substitution rates at a lower limit of detection of 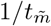 (Figure S11). This limit of detection corresponds to the lowest possible non-zero rate that can be measured for mutations where 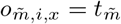. This clipping only affected a small number of synonymous mutations with much lower rates than average for a given evolutionary opportunity, such as the synonymous mutations with highly deleterious fitness effects from Figure 4.

When computing mutation rates, we excluded synonymous mutations in the following regions. From the start of our analysis, we excluded them in overlapping reading frames (ORFs) in the PA, MP, and NS segments, unless they were synonymous in both frames. For PA, the overlap was between the PA and X [58] (sites 572-760 in our PA reference positive-sense coding sequence) ORFs. For MP, it was between the M1 and M2 ORFs. For NS, it was between the NS1 and NEP ORFs. When analyzing fitness effects of the remaining synonymous mutations, we identified two other regions where synonymous mutations were under strong purifying selection, which we ultimately excluded when computing mutation rates, as well. We excluded synonymous mutations in regions at segment termini with known packaging signals, with region boundaries taken from Li et al. [39]. We also excluded synonymous mutations near exon/intron boundaries involved in splicing of the M2 and NEP genes (for each such boundary, we excluded all sites in a window of 10 sites centered on the boundary). See https://github.com/matsengrp/flu-mut-rates/blob/main/data for:

- `packaging signal boundaries.csv:` a file giving boundaries of exclusion for packaging signals
- `splice site boundaries.csv:` a file giving locations of exon/intron boundaries, determined using annotations from reference sequences used to build trees (Table S4).

See https://github.com/matsengrp/flu-mut-rates/blob/main/results for files with rates, including:

- `genome_wide_rates.csv:` genome-wide substitution rates 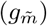
- `motif_level_genome_wide_rates.csv:` genome-wide motif-specific sub-stitution rates 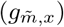
- `site_specific_substitution_rates.csv`: site-specific substitution rates 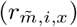
- `evo_opp_thresholds.csv:` 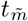 values for each mutation type

The repository’s `README.md` file describes each of the above files in more detail.

### Fitting a neutral mutation model

From the empirical rates 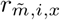, restricted to synonymous mutations, we estimate neutral rates of mutation with a log-linear model. For each mutation type, we collect the logarithm of the synonymous substitution rate and sequence context at each site. We separately fit a model for each mutation type, with data from all sequence contexts appearing with the mutation type, so that each model takes the form

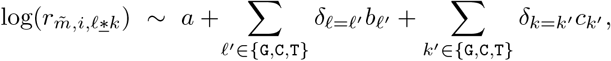

where *δ*_(•)_ is the indicator function for the given condition. That is to say, each log-rate is determined by a constant term, a term for the left neighboring nucleotide, and a term for the right neighboring nucleotide (7 terms, as opposed to 16). The nucleotide A is taken as the reference category, meaning the contribution is in the constant term, rather than in the sums. The terms are selected to minimize mean squared error, as the logarithm of the rates is assumed normal, with L2-regularization (ridge regression) to limit overfitting. We use the standard closed-form estimator

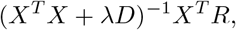

where *X* is the matrix encoding the local sequence context at each site (each context is recorded as a vector of length 7), *λ* is the L2-penalty term (we use 0.1), *D* is the diagonal matrix with diagonal [0, 1, 1, …, 1] (the first term is zero because the penalty should not apply to the constant term), and *R* is the vector of the empirical log-rate at each site. This results in the estimated neutral rates for each mutation type and sequence context.

As rates may vary from one genomic segment to another, we also fit models with a term for the genomic segment. Specifically, these models take the form

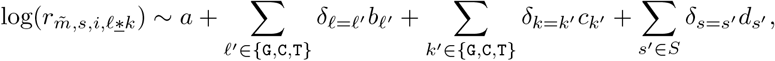

where *s* is the segment in which the mutation occurred and *S* is the set of segments {PB2,PA,HA,NP,NA,MP,NS} (note either *s* ∈ *S* or *s* is PB1, which is taken as the reference category). Fitting such a model is the same as before. These models account for substitution rates following similar patterns among segments, but at different orders of magnitude.

See https://github.com/matsengrp/flu-mut-rates for files with model-predicted substitution rates.

### Estimating mutational fitness effects

The *Results* section describes our basic approach for doing this. Here, we expand using more detailed notation that mirrors the rest of the *Methods* section.

We defined the actual count of a nucleotide-level mutation of type 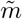 at site *i* in context *x* as equal to 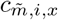 described above 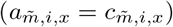. We computed an expected count for that mutation as 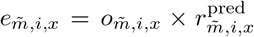, where 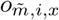 is the mutation’s evolutionary opportunity and 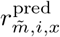 is the predicted rate of a mutation according to the neutral mutation model described above that models both local sequence context and genomic segment. For C→G and G→C mutations, which are not included in the neutral model, we predicted a mutation’s rate as the genome-wide rate of mutations in that context 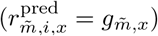 for the 12 of 16 contexts with 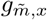 estimates, or as the context-independent genome-wide rate 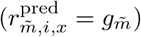 for the remaining four contexts not present in the data (due to synonymous mutations only being possible in certain contexts).

When computing fitness effects of nucleotide-level mutations, we aggregated counts across sequence contexts. Specifically, for a mutation 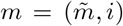, we defined aggregated actual and expected counts as 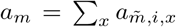, and 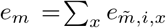 we computed the effect of *m* as:

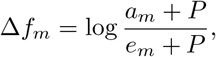

where *P* is a pseudocount of 0.5 to avoid division by zero, matching the notation used in the *Results*. The equation uses the natural log.

When computing fitness effects of amino-acid mutations, we aggregated counts across sequence contexts and nucleotide-level mutations encoding the same amino-acid change. Analogously to the nucleotide case, we use 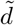 to de-note an amino-acid mutation type and *n* to denote a codon site, and we write 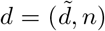 for an amino-acid mutation at a specific codon site. Specifically, we computed the effect of amino-acid mutation *d* as:

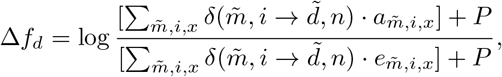

where 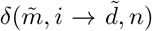 is an indicator variable equal to 1 if nucleotide mutation 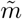 at site *i* results in amino-acid mutation 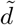 at codon site *n*.

We only computed fitness effects for mutations with at least ten actual or expected counts (after aggregation steps described above).

In overlapping reading frames in the PA, MP, and NS segments, we classified mutations as follows. We only classified a mutation as being synonymous if it was synonymous in both reading frames. We classified mutations as nonsense if they were nonsense in at least one frame. Otherwise, we classified mutations as nonsynonymous if they were nonsynonymous in at least one frame.

See https://matsen.group/flu-mut-rates for interactive heatmaps of fitness effects of mutations to reference sequences used to build each tree. We made the heatmaps using code adapted from the `polyclonal` software package [59]. See https://github.com/matsengrp/flu-mut-rates/blob/main/ results for files with fitness effects, including:

- `nt fitness effects.csv:` fitness effects of nucleotide-level mutations
- `aa fitness effects.csv:` fitness effects of amino-acid mutations

## Acknowledgments

We gratefully acknowledge all data contributors, i.e., the authors and their originating laboratories responsible for obtaining the specimens, and their submitting laboratories for generating the genetic sequence and metadata and sharing via the GISAID Initiative, on which this research is based (Table S1). We thank Mary Barker and Ognian Milanov for assistance with larch. We thank Richard Neher and Russell Corbett-Detig for useful discussions.

## Funding

KJ and CTB were supported by the Fred Hutch REACH (Research Equity Advancement for Cancer with HBCUs) program, a collaboration between the Fred Hutch and Historically Black Colleges and Universities, funded by the Fred Hutch and generous contributions from private donors (The Ticknor Family and The Havens Family). This research was supported in part by grant no. R01 AI146028 from the NIH. FAM and JDB are investigators of the Howard Hughes Medical Institute. Scientific Computing Infrastructure at Fred Hutch funded by ORIP grant S10OD028685.

## Disclosures

JDB consults for Apriori Bio, Invivyd, the Vaccine Company, Pfizer, GSK, and Merck.

## Notes

https://github.com/matsengrp/flu-usher

https://github.com/matsengrp/flu-mut-rates

