## Supplemental Information for "Phylogenetically estimated neutral rates and fitness effects of mutations to influenza proteins"

### Supplementary Information for: Phylogenetically estimated neutral rates and fitness effects of mutations to influenza proteins

Hugh K. Haddox, Angie S. Hinrichs, Chris Jennings-Shaffer, Karrington Johnson, Chelsea T. Benton, Jared G. Galloway, Jesse D. Bloom, Frederick A. Matsen IV

May 15, 2026

### Tables

| Subtype | Maximum submission date | EPISET ID | DOI |
|---|---|---|---|
| H1N1 | 2025-12-01 | EPI_SET_260123kw | 10.55876/gis8.260123kw |
| H3N2 | 2025-12-01 | EPI_SET_260123av | 10.55876/gis8.260123av |
| H5N1 | 2025-05-30 | EPI_SET_260123qh | 10.55876/gis8.260123qh |
| H7N9 | 2025-05-19 | EPI_SET_260123ou | 10.55876/gis8.260123ou |
| H9N2 | 2025-06-03 | EPI_SET_260123ek | 10.55876/gis8.260123ek |
| H1NX | 2025-07-15 | EPI_SET_260123cz | 10.55876/gis8.260123cz |
| H3NX | 2025-07-15 | EPI_SET_260123pn | 10.55876/gis8.260123pn |
| H5NX | 2025-07-15 | EPI_SET_260123xb | 10.55876/gis8.260123xb |
| H7NX | 2025-07-15 | EPI_SET_260123mt | 10.55876/gis8.260123mt |
| H9NX | 2025-07-15 | EPI_SET_260123qa | 10.55876/gis8.260123qa |
| HXNX | 2025-07-15 | EPI_SET_260123om | 10.55876/gis8.260123om |

**Table S1: Maximum submission dates and EPISET IDs for influenza sequences downloaded from GISAID.** Rows show data for a given subtype. If the H or N subtype specification is an "X" (e.g., H1NX), then that includes all subtypes for that H or N, discounting subtypes previously listed in the table (e.g., H1NX would include all subtypes with the H1 specification, except for H1N1, which is listed previously in the table). tab:episet

| Host | Number of sequences |
|---|---:|
| human | 422,767 |
| avian | 85,019 |
| swine | 28,594 |
| other | 10,154 |
| bovine | 3,969 |

**Table S2: Distribution of sequences by host group.** tab:hosts

| Subtype | Number of sequences |
|---|---:|
| H3N2 | 226,173 |
| H1N1 | 216,388 |
| H5N1 | 38,663 |
| H9N2 | 18,814 |
| H1N2 | 9,723 |
| H3N8 | 5,217 |
| H5N8 | 4,785 |
| H7N9 | 3,216 |
| H5N6 | 3,211 |
| H4N6 | 2,772 |

**Table S3: Distribution of sequences by subtype for the top ten most common subtypes.** tab:subtypes

| Segment | Subtype | Reference Accession | Notes |
|---|---|---|---|
| HA | H1 | LN867357.1 | A/England/414/2010 |
| HA | H3 | KF874500.2 | A/Aichi/2/1968 |
| HA | H5 | NC_007362.1 | A/goose/Guangdong/1/1996 |
| HA | H7 | NC_026425.1 | A/Shanghai/02/2013 |
| HA | H9 | NC_004908.1 | A/Hong Kong/1073/99 |
| NA | N1 | NC_002018.1 | A/Puerto Rico/8/1934 |
| NA | N2 | CY121119.1 | A/Aichi/2/1968 |
| NA | N6 | KP732684.1 | A/duck/Eastern China/S0322/2014 |
| NA | N8 | L06575.1 | A/Duck/Memphis/928/74 |
| NA | N9 | NC_026429.1 | A/Shanghai/02/2013 |
| PB2 | all | CY121124.1 | A/Aichi/2/1968 (H3N2) |
| PB1 | all | CY121123.1 | A/Aichi/2/1968 (H3N2) |
| PA | all | CY121122.1 | A/Aichi/2/1968 (H3N2) |
| NP | all | CY121120.1 | A/Aichi/2/1968 (H3N2) |
| MP | all | CY121118.1 | A/Aichi/2/1968 (H3N2) |
| NS | all | CY121121.1 | A/Aichi/2/1968 (H3N2) |

**Table S4:** Reference accession numbers for each segment-subtype combination, from NCBI (`https://www.ncbi.nlm.nih.gov/`)

tab:reference˙accessions

| Segment | Subtype | Root Node (EPI_ISL) |
|---|---|---|
| HA | H1 | EPI_ISL_5897 |
| HA | H3 | EPI_ISL_18852459 |
| HA | H5 | EPI_ISL_5886 |
| HA | H7 | EPI_ISL_10304 |
| HA | H9 | EPI_ISL_3214 |
| NA | N1 | EPI_ISL_5878 |
| NA | N2 | EPI_ISL_8017 |
| NA | N9 | EPI_ISL_87301 |
| PB2 | all | EPI_ISL_5886 |
| PB1 | all | EPI_ISL_5886 |
| PA | all | EPI_ISL_5886 |
| NP | all | EPI_ISL_5886 |
| MP | all | EPI_ISL_5886 |
| NS | all | EPI_ISL_5886 |

**Table S5: Sequences used for tree rerooting.** Each sequence comes from an avian isolate with a relatively early collection date. The N6 and N8 trees were not rerooted.

tab:reroot˙sequences

### Figures

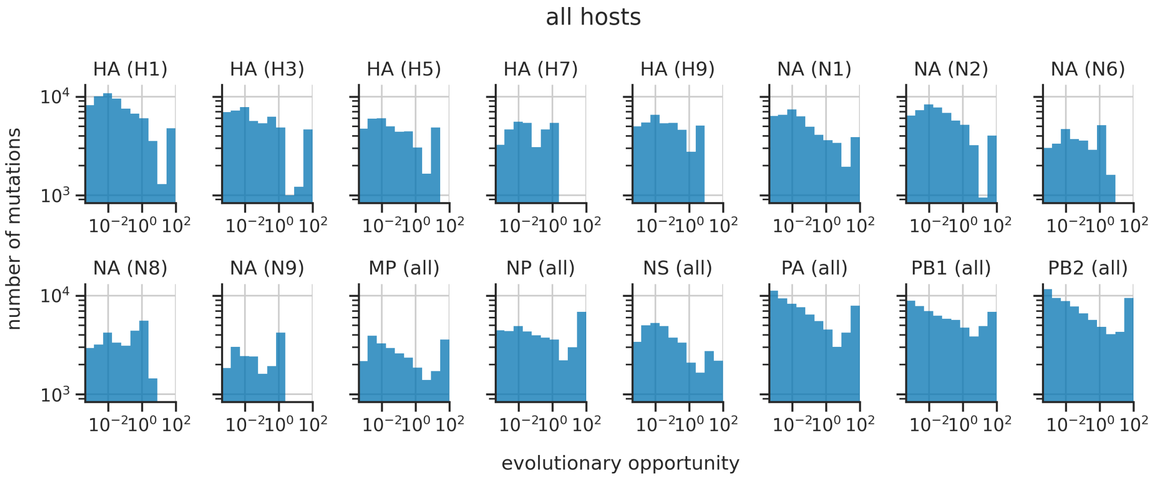


**Figure S1: Distribution of the evolutionary opportunity of mutations.** Each histogram shows data from a tree for a specific segment and subtype, with leaf sequences from all hosts. Histograms bin mutations by their evolutionary opportunity ($o_{\tilde{m},i,x}$ values defined in the *Methods* section). The distribution is over the set of all possible single-nucleotide mutations to parent nodes of branches on the tree. fig:evo˙opp

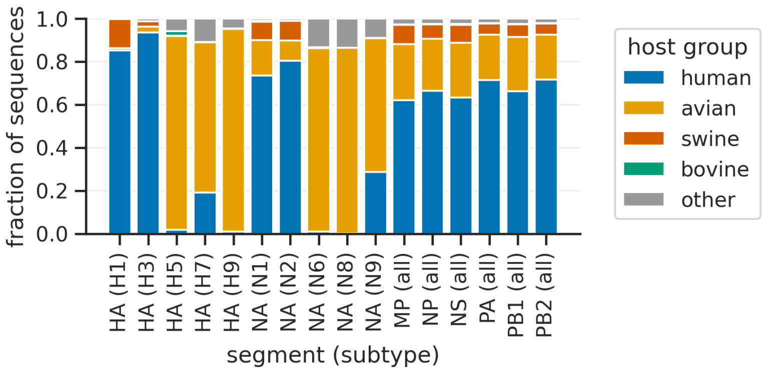


**Figure S2: Fraction of sequences per tree from a given host group.** fig:frac˙seqs˙by˙host

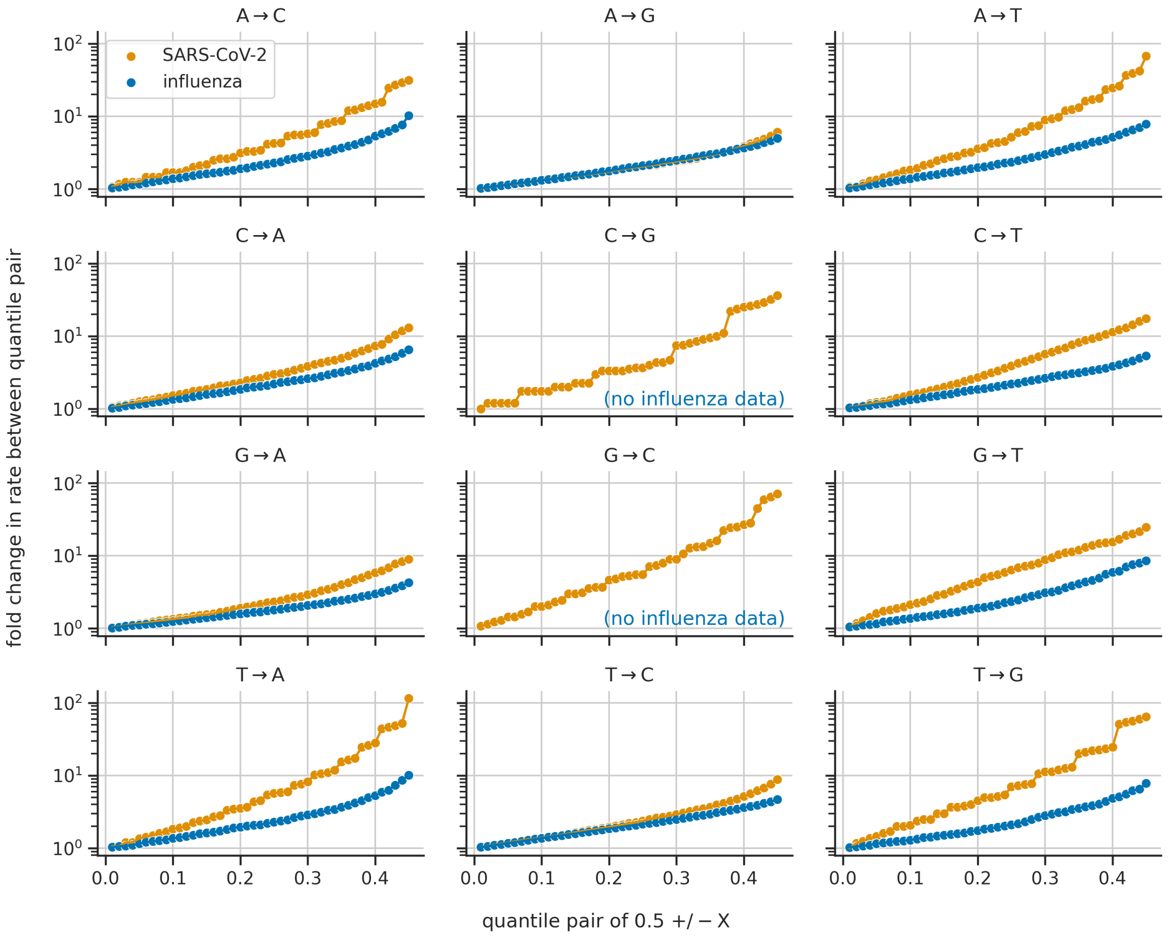


**Figure S3: Site-specific synonymous substitution rates are less variable for influenza than for SARS-CoV-2.** This figure examines distributions of site-specific synonymous substitution rates associated with a specific mutation type and virus. For a given mutation type and virus, we quantified the distribution's variability by computing fold changes in rates between a series of upper and lower quantile pairs. We considered pairs of $0.5 + / - X$, where $X$ is the value plotted on the x axis. For instance, $X = 0.1$ corresponds to a lower quantile of 0.4 and an upper quantile of 0.6. The y axis shows the fold change in rates between quantile pairs, computed as the upper quantile's rate divided by the lower quantile's rate. Blue dots show results for influenza, while orange dots show results for SARS-CoV-2. Fold changes tend to be lower for influenza, indicating reduced variability in rates. The SARS-CoV-2 data is from Haddox et al. [1]. fig:rate˙variation˙comparison

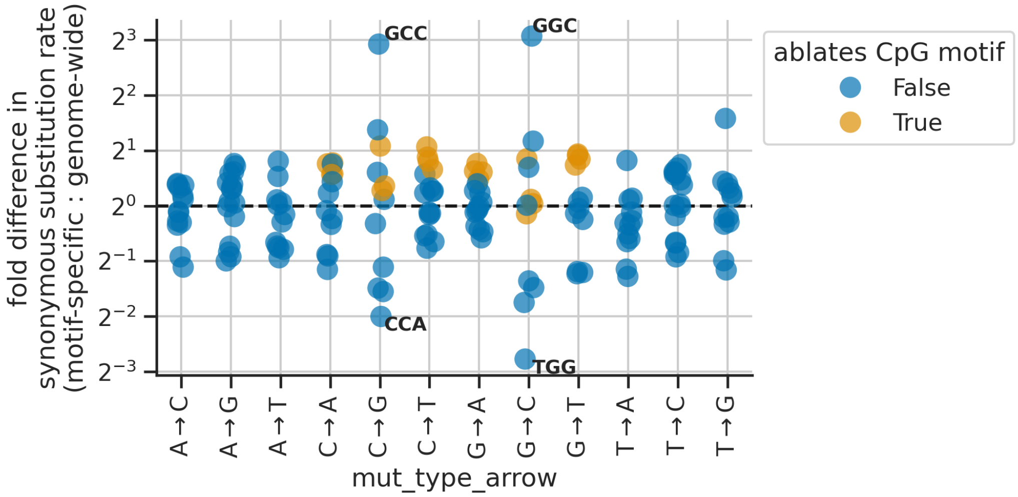


**Figure S4: Variation in synonymous substitution rates between 3-mer motifs.** For each mutation type, the plot shows the fold difference between motif-specific synonymous substitution rates compared to the genome-wide synonymous substitution rate for that type. Each dot represents a different 3-mer motif. For most mutation types, motif-specific rates span a range from about two-fold lower to about two-fold higher than the genome-wide rate. A few motif-specific rates lie outside of this range, with the most extreme outliers labeled by their motif. Dots are colored based on whether mutating the central position of the motif disrupts a `CG` dinucleotide in the genome, which is a target of ZAP. For a given mutation type, mutations that disrupt `CG` motifs often have higher rates than mutations that do not (orange dots tend to have higher rates than blue dots in the same column). Motif-specific and genome-wide rates are computed as averages across the genome, as described in detail in the *Methods* section (see definitions of $g_{\tilde{m},x}$ and $g_{\tilde{m}}$). fig:motif˙rate˙variation

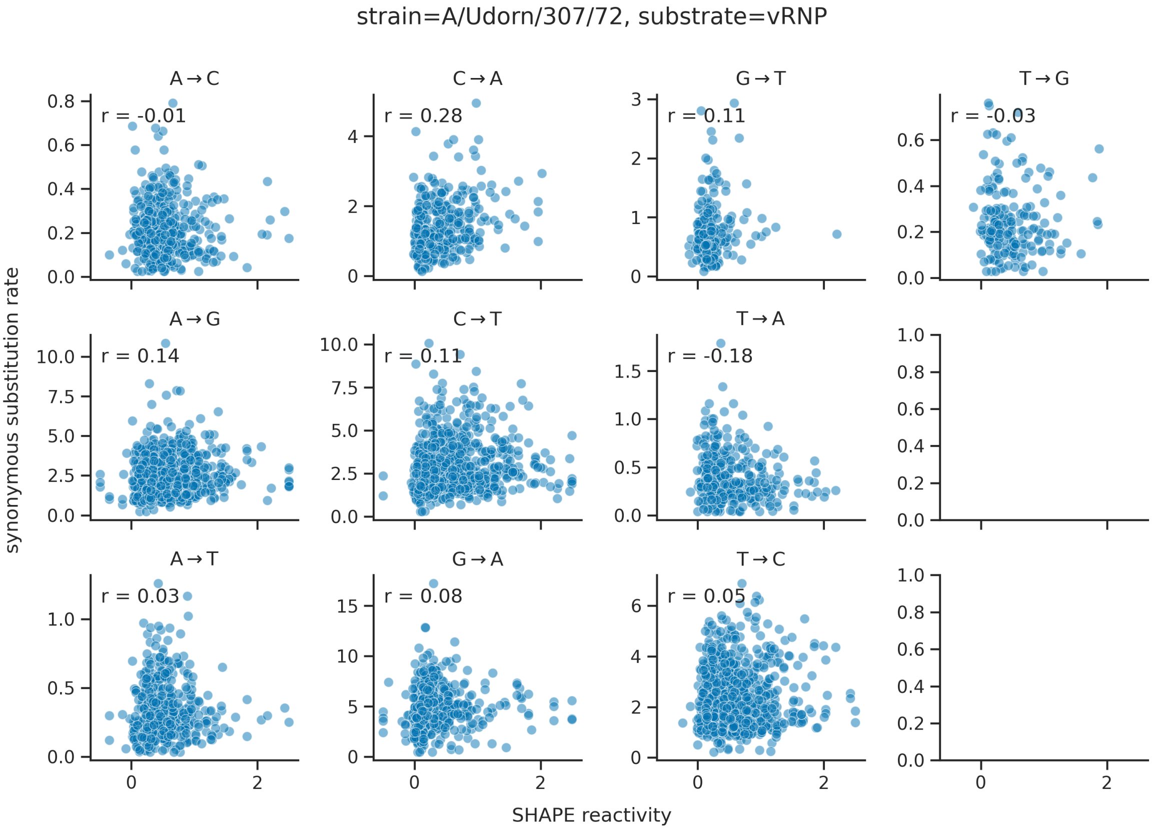


**Figure S5: Site-specific synonymous substitution rates do not strongly correlate with experimental SHAPE-Map data quantifying RNA base pairing.** For each mutation type, we plotted the correlation between site-specific synonymous substitution rates and site-specific SHAPE reactivity values measured by Dadonaite et al. [2]. In each plot, each dot corresponds to a specific site in the genome, and $r$ reports the Pearson correlation coefficient. The correlations are low. We excluded sites in the HA and NA segments, since these segments are highly variable, focusing our analysis on the remaining, more conserved segments. We show reactivity values measured in context of the A/Udorn/307/72 strain and a vRNP substrate, but the results were consistent between strains and substrates, as shown by similar plots in `https://github.com/matsengrp/flu-mut-rates/blob/main/notebooks/analyze_site_specific_rates.ipynb`. This figure does not show data for C→G and G→C mutations, since we could not estimate site-specific rates for these mutation types. fig:rna ss

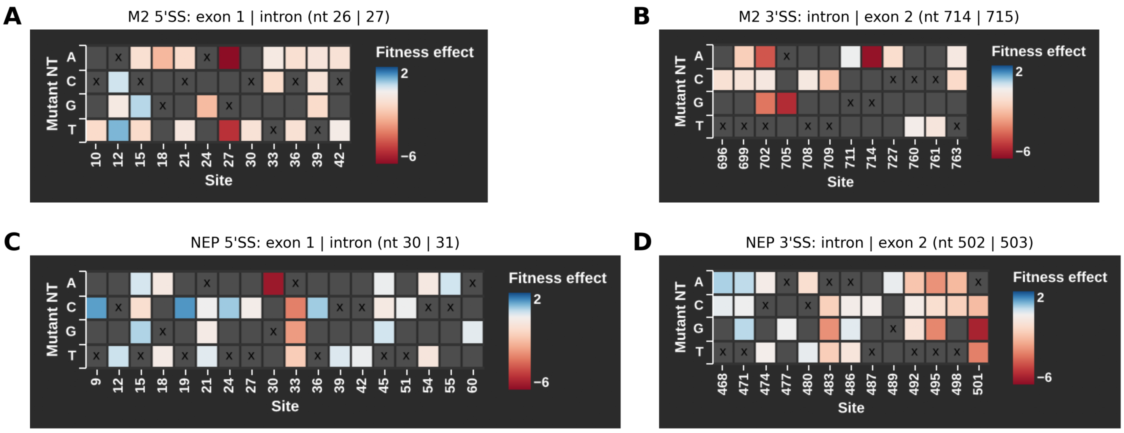


**Figure S6: Fitness effects of nucleotide mutations near exon/intron boundaries involved in splicing of M2 and NEP.** Each panel shows a heatmap of fitness effects of all synonymous mutations with estimated fitness effects near a given boundary. Site numbers are not sequential, since we only show data for sites with estimated synonymous-mutation effects. **(A)** This panel shows sites in the region of the 5′ splice site of the M2 coding sequence, with the boundary located at the end of exon 1 (site 26) and the beginning of the subsequent intron (site 27), with sites numbered in context of the positive-sense MP segment's coding sequence. **(B)** This panel is similar to panel A, but shows sites in the region of the 3′ splice site of the M2 coding sequence, with the boundary located at the end of the intron (site 714) and the beginning of exon 2 (site 715). **(C)** and **(D)** These panels are similar to panels A and B, but show data for 5′ and 3′ splice sites of the NEP coding sequence (see titles for the location of boundary sites). In panels A-C, there is at least one synonymous mutation with an estimated fitness effect at a boundary site (e.g., site 27 in panel A). All of these effects are deleterious. In panel D, the closest mutations with estimated effects are at site 501, which is one site away from the boundary. Mutations are deleterious at this site, too. fig:ss˙heatmaps

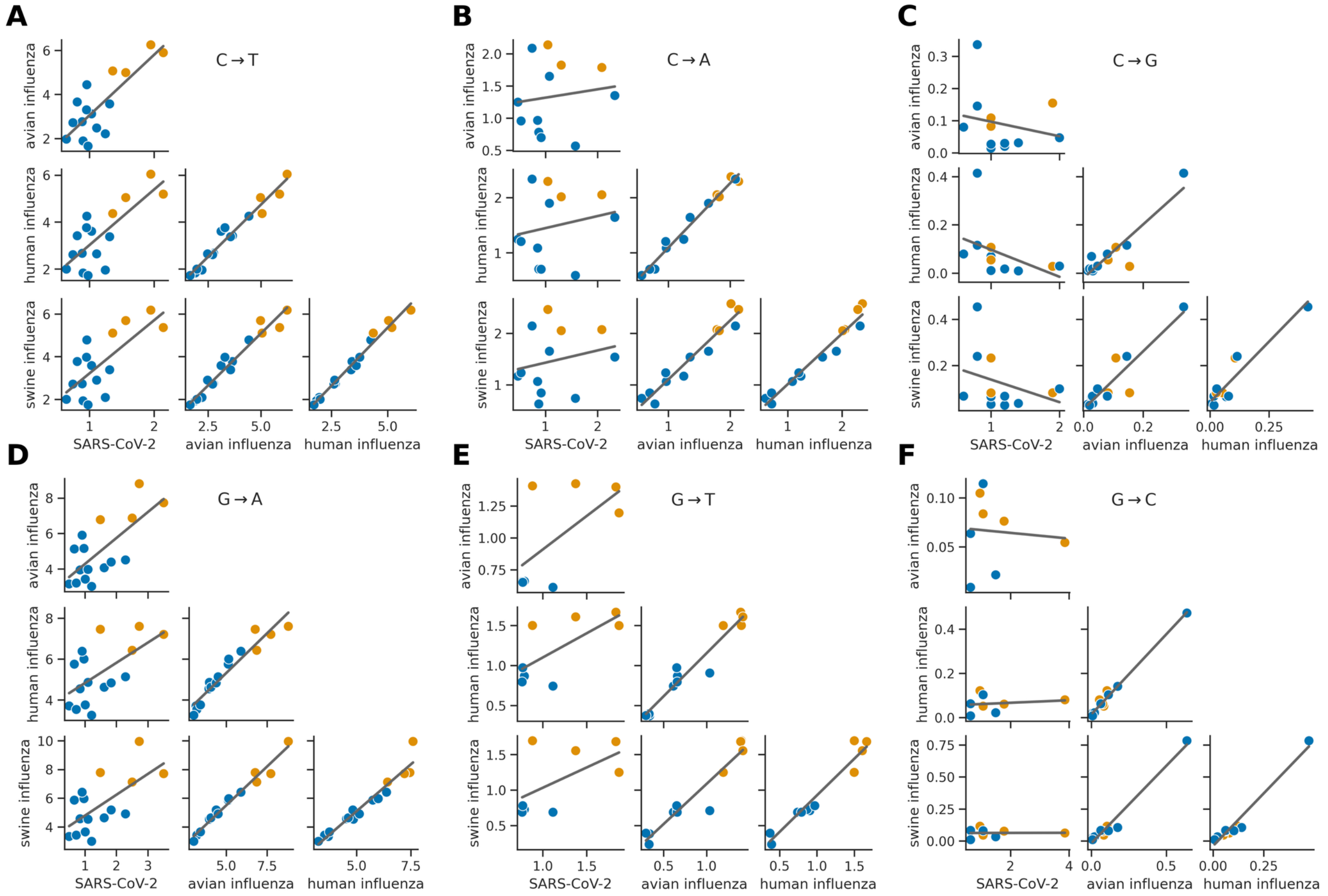


**Figure S7: The elevation in substitution rates of synonymous mutations that ablate `CG` motifs is consistent between hosts, and is sometimes consistent between influenza and SARS-CoV-2.** Figure S4 shows that substitution rates of synonymous mutations that ablate `CG` dinucleotide motifs are often higher compared to mutations that do not, for a given mutation type. That figure shows host-agnostic substitution rates. This figure shows the correlation of host-specific substitution rates between human influenza, swine influenza, avian influenza, or SARS-CoV-2, with one panel for each mutation type going away from a `C` nucleotide (panels A-C) or a `G` nucleotide (panels D-F). As in Figure S4, each dot corresponds to a different 3-mer motif and its value corresponds to the substitution rate of synonymous mutations in context of that motif for a given virus. Orange dots indicate 3-mer motifs where mutations to the central nucleotide ablate a `CG` dinucleotide motif, while blue dots show other motifs. The estimated motif-specific rates are highly correlated between human, swine, and avian influenza, including for mutation types where rates are elevated for mutations that ablate `CG` dinucleotide motifs, such as for C→T (the orange dots have higher rates than the blue dots across hosts). For C→T and G→A, this pattern is also consistent between influenza and SARS-CoV-2.
fig:motif specific rates pairplots

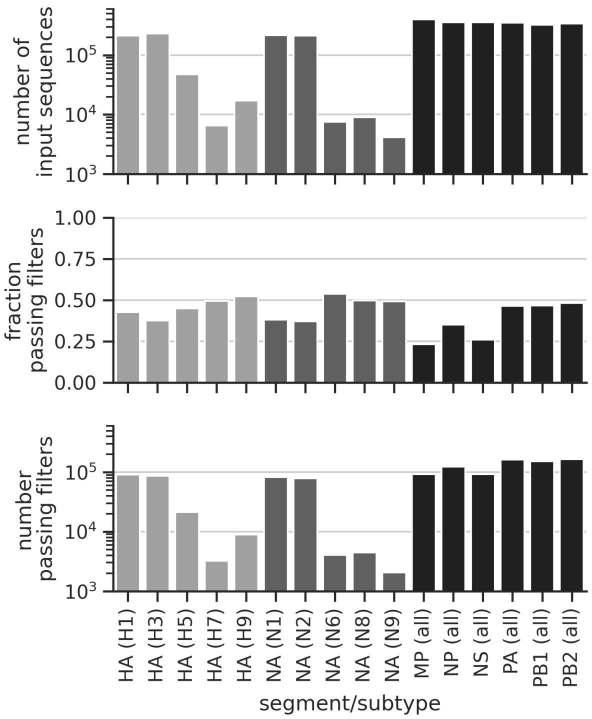


**Figure S8: Number of sequences for a given segment and subtype combination before and after filtering.** For the HA and NA segments, we made separate trees for each subtype. For the remaining segments, we made a single tree with sequences from all subtypes. The top plot shows the raw number of input sequences for a given combination. The middle plot shows the fraction of sequences that were filtered out upon the alignment and filtering steps described in the main text. Most sequences were filtered out because they were duplicates of other sequences. The bottom plot shows the final number of sequences included in each tree. fig:alignment filtering stats

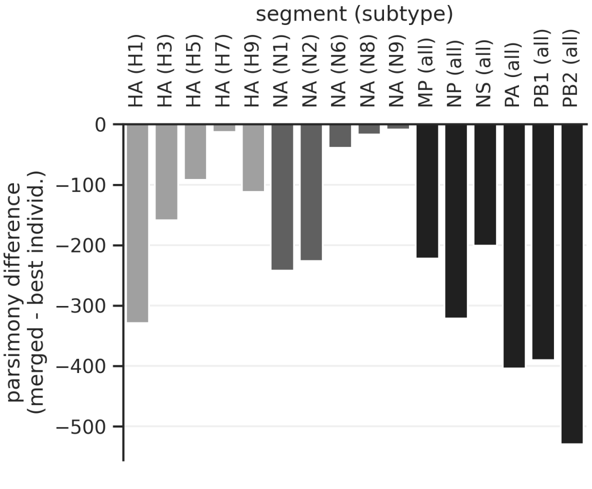


**Figure S9: Improvement in parsimony score between the tree extracted from the merged `larch` DAG and the best of the ten input trees.** Each bar shows results for a given segment and subtype combination put through the tree-building pipeline (e.g., HA (H1) shows data for the HA segment and H1 subtype). Lower parsimony scores are better, so the negative values on the y axis indicate that the tree extracted from the merged DAG has a better score. The trees with larger improvements (more negative values) tend to be the trees with more sequences (Figure S8), which makes sense given that parsimony scores scale with tree size.

fig:delta`parsimony

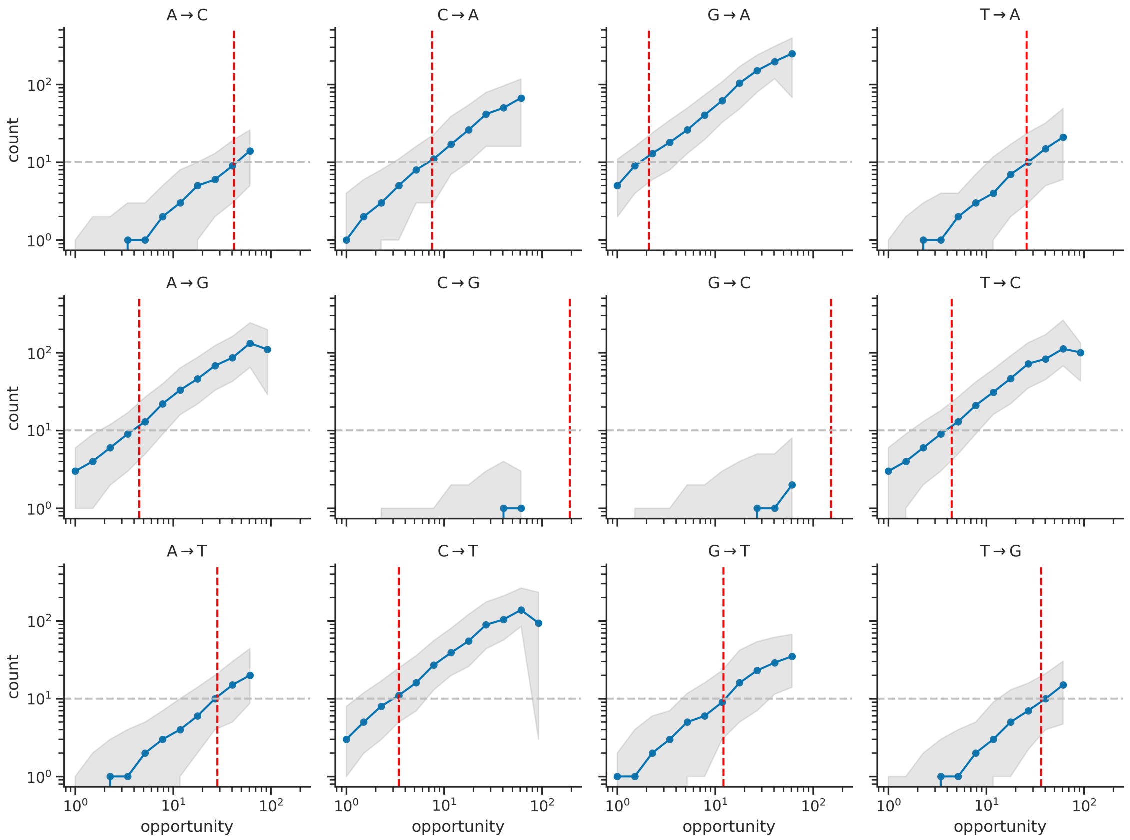


**Figure S10: Distribution of mutation counts as a function of evolutionary opportunity.** Each plot shows data for a given mutation type. The x axis bins mutations by evolutionary opportunity, using bins that are evenly spaced in log space (only showing data for mutations with opportunity values ≥1.0). The y axis shows the distribution of mutation counts in that bin, with the blue line showing the median and the shaded region showing the range between the 10th and 90th quantiles. Each blue dot shows the median for a given bin, where the dot's value on the x axis corresponds to the bin's minimum value (all values between a given dot and up to but not including the dot to its right are in a single bin). Mutation counts scale linearly with evolutionary opportunity. Vertical dashed red lines show the evolutionary opportunity that is expected to result in an average of ten counts per site ($t_{\tilde{m}}$; see *Methods*). Indeed, the blue line connecting the dots is close to a value of ten where it intersects the red line. And the shaded region indicates that most mutations have at least one, and often several counts at the red line. No G→C or C→G mutations had sufficient evolutionary opportunity to reach an expected value of ten counts per site (red line), which is confirmed by the observed counts distribution (blue line and shaded region). fig:counts˙by˙opp

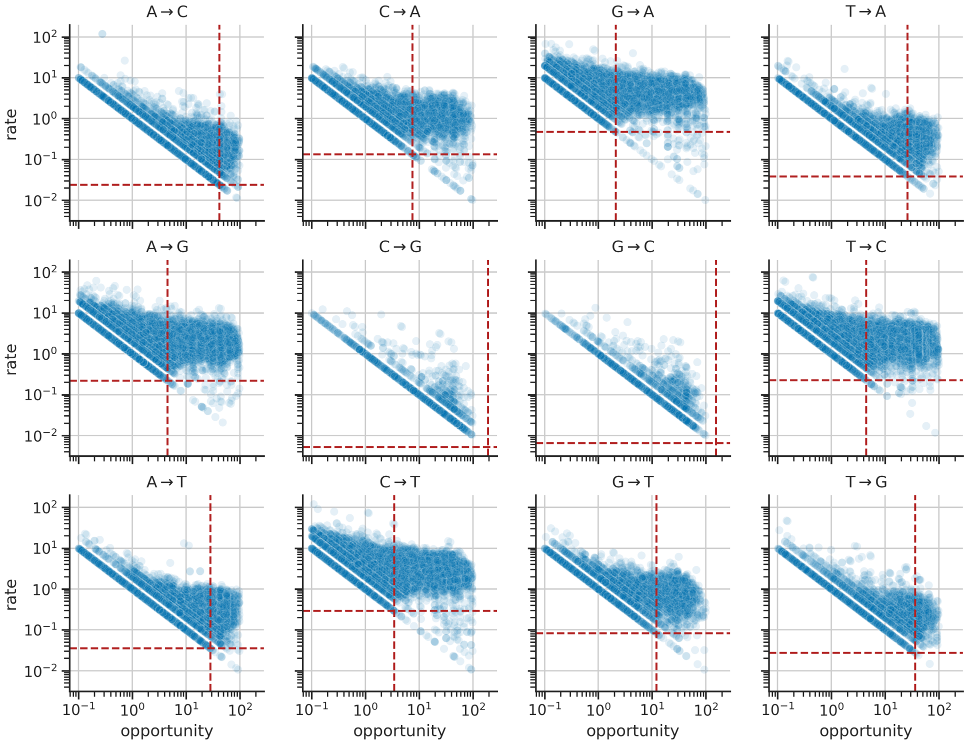


**Figure S11: Distribution of substitution rates as a function of evolutionary opportunity.** Each plot shows data for a given mutation type. Each dot corresponds to a specific synonymous mutation at a specific site in the genome. The x axis shows the mutation's evolutionary opportunity ($o_{\tilde{m},i,x}$), while the y axis shows its rate ($r_{\tilde{m},i,x}$). In each plot, the diagonal floor of points corresponds to mutations with a count of 1.0, such that $r_{\tilde{m},i,x} = 1/o_{\tilde{m},i,x}$. Vertical dashed red lines show the evolutionary opportunity that is expected to result in an average of ten counts per site ($t_{\tilde{m}}$; see *Methods*). We only analyze site-specific substitution rates for mutations with $o_{\tilde{m},i,x}$ values greater than or equal to this threshold. Horizontal dashed red lines show the lower limit of detection of substitution rates for mutations with $o_{\tilde{m},i,x}$ values equal to the threshold, where the limit of detection is the floor described above ($r_{\tilde{m},i,x} = 1/o_{\tilde{m},i,x}$). We clipped site-specific substitution rates at this lower limit of detection.

fig:rates˙by˙opp
